# Genomics of Natural Populations: Evolutionary Forces that Establish and Maintain Gene Arrangements in *Drosophila pseudoosbscura*

**DOI:** 10.1101/177204

**Authors:** Zachary L. Fuller, Gwilym D. Haynes, Stephen Richards, Stephen W. Schaeffer

## Abstract

The evolution of complex traits in heterogeneous environments may shape the order of genes within chromosomes. *Drosophila pseudoobscura* has a rich gene arrangement polymorphism that allows one to test evolutionary genetic hypotheses about how chromosomal inversions are established in populations. *D. pseudoobscura* has >30 gene arrangements on a single chromosome that were generated through a series of overlapping inversion mutations with > 10 inversions with appreciable frequencies and wide geographic distributions. This study analyzes the genomic sequences of 54 strains of *Drosophila pseudoobscura* that carry one of six different chromosomal arrangements to test whether (1) genetic drift, (2) hitchhiking with an adaptive allele, (3) direct effects of inversions to create gene disruptions caused by breakpoints, or (4) indirect effects of inversions in limiting the formation of recombinant gametes are responsible for the establishment of new gene arrangements. We found that the inversion events do not disrupt the structure of protein coding genes at the breakpoints. Population genetic analyses of 2,669 protein coding genes identified 277 outlier loci harboring elevated frequencies of arrangement-specific derived alleles. Significant linkage disequilibrium occurs among distant loci interspersed between regions with low levels of association indicating that distant allelic combinations are held together despite shared polymorphism among arrangements. Outlier genes showing evidence of genetic differentiation between arrangements are enriched for sensory perception and detoxification genes. The data presented here support the indirect effect of inversion hypothesis where chromosomal inversions are favored because they maintain linked associations among multi-locus allelic combinations among different arrangements.

## Introduction

Chromosome structure and genome organization have been fluid over evolutionary history. An important force that may shape how genes are distributed across the genome and their order on chromosomes may be the evolution of complex traits. For single loci, natural selection is more efficient and effective in the presence of recombination because beneficial mutations can be dissociated from linked deleterious alleles (Becks & Agrawal, 2010, 2012; McDonald, Rice, & Desai, 2016; Otto & Lenormand, 2002) On the other hand, for traits controlled by multiple loci, recombination can break apart favorable combinations of alleles just as easily as it can bring them together. Therefore, the architecture of the genome may be shaped by mechanisms that modulate recombination rates such as fusing chromosomes together (Dobigny, Britton-Davidian, & Robinson, 2015; McAllister, Sheeley, Mena, Evans, & Schlotterer, 2008) or generating karyotypic variation among closely related species (Carbone et al., 2014; Nachman, Boyer, Searle, & Aquadro, 1994; O’Neill, Eldridge, & Metcalfe, 2004; O’Neill et al., 1999).

Genes within chromosome can be shuffled through meiotic recombination, which can be reduced by the presence of chromosomal inversions (Sturtevant, 1926). Inversions reduce recombination in rearrangement heterozygotes because unbalanced gametes form as a result crossing over within the inverted segments. In humans, inversions are discovered in patients with disease symptoms and are found to have chromosomal breaks in vital genes (Pettenati et al., 1995). With the advent of genomic sequencing, non-disease causing inversions have been discovered to be segregating in human populations (Stefansson et al., 2005).

Inversions are segregating in a wide variety of taxa (Coluzzi, Sabatini, Petrarca, & Di Deco, 1979; Engelbrecht, Taylor, Daniels, & Rambau, 2011; Kupper et al., 2016; Lee, Fishman, Kelly, & Willis, 2016; Pyhäjärvi, Hufford, Mezmouk, & Ross-Ibarra, 2013; Silva et al., 2014; Diether Sperlich & Pfriem, 1986; Stefansson et al., 2005; Zinzow-Kramer et al., 2015) and are assumed to lead to the wealth of fixed chromosomal rearrangements observed between species (W. W. Anderson, Ayala, & Michod, 1977; Carson, 1992; Coluzzi, Sabatini, della Torre, Di Deco, & Petrarca, 2002; Fontaine et al., 2015; Levitan, 1992; McGraw, Davis, Young, & Thomas, 2011; Nishikawa et al., 2015; Wasserman, 1982). Indirect evidence has suggested that natural selection modulates the frequencies of gene arrangements because many show clinal variation across environmental gradients (Balanya et al., 2003; Cheng et al., 2012; Dobzhansky, 1944; Martin Kapun, Fabian, Goudet, & Flatt, 2016; M. Kapun, Schmidt, Durmaz, Schmidt, & Flatt, 2016; M. Kapun, van Schalkwyk, McAllister, Flatt, & Schlotterer, 2014; Knibb, 1982; Knibb, Oakeshott, & Gibson, 1981; Mettler, Voelker, & Mukai, 1977) including independent parallel clines observed in northern and southern hemispheres, as well across different continents (A. R. Anderson, Hoffmann, McKechnie, Umina, & Weeks, 2005; Calboli, Kennington, & Partridge, 2003; Knibb, 1982; Kolaczkowski, Kern, Holloway, & Begun, 2011; M. Santos et al., 2005). For example, natural selection acting on inversion polymorphism in *D. buzzattii* is thought to maintain the genetic architecture underlying thermal adaptation across an environmental gradient (Soto et al., 2010). Furthermore, environmental selection is hypothesized to shape the distribution of chromosomal arrangements in the malaria vector *Anopheles funestus* (D. Ayala et al., 2011) and inversions are implicated in clinal adaptation in other *Anopheles* species (Diego Ayala et al., 2017). Despite their pervasiveness both as polymorphisms within species and fixed differences between species, the mechanisms that establish and maintain different gene arrangements in populations are unclear.

Four broad classes of hypotheses have been proposed to explain how inversions are established in populations: (1) mutation and genetic drift, (2) hitchhiking with an adaptive mutation, (3) direct effects of the inversion mutation, and (4) indirect effects of suppressed recombination (Hoffmann & Rieseberg, 2008; Kirkpatrick & Barton, 2006). A new gene arrangement can be established by drift in small isolated populations subject to high rates of extinction and recolonization (Lande, 1984). An inversion could also rise to high frequency if it captures an adaptive allele that is sweeping through the population (Maynard Smith & Haigh, 1974). Sperlich (1959) suggested that inversions could directly create selectable variation by altering structure or expression of genes flanking breakpoints (Fuller, Haynes, Richards, & Schaeffer, 2016). Another direct effect of inversions might be in their potential to form segregation distorters (Novitski, 1951). Inversions may also be established through the maintenance of linked associations of alleles because of their indirect effect of reducing recombinant gametes (Sturtevant & Beadle, 1936) either because the chromosome is free of deleterious recessive alleles (Nei, Kojima, & Schaffer, 1967; Ohta, 1971), the chromosome has epistatic combinations of alleles (Charlesworth & Charlesworth, 1973; Dobzhansky, 1950) or the chromosome has locally adapted alleles (Kirkpatrick & Barton, 2006).

*Drosophila pseudoobscura* has been used as a model system to investigate inversions in natural populations with > 30 different gene arrangements generated through single inversion events (Dobzhansky & Sturtevant, 1938; Powell, 1992). The polymorphism is estimated to be 1.4-1.7 million years old (Aquadro, Weaver, Schaeffer, & Anderson, 1991; A. G. Wallace, Detweiler, & Schaeffer, 2011). Several lines of direct and indirect evidence suggest that selection helps establish and maintain the different gene arrangements. Five gene arrangements are frequent, widely distributed, and form clines across the southwestern United States where frequency shifts coincide with changes in major physiographic provinces (Dobzhansky, 1944; Lobeck, 1948; Schaeffer, 2008) (Figure 1).

**Figure 1.**
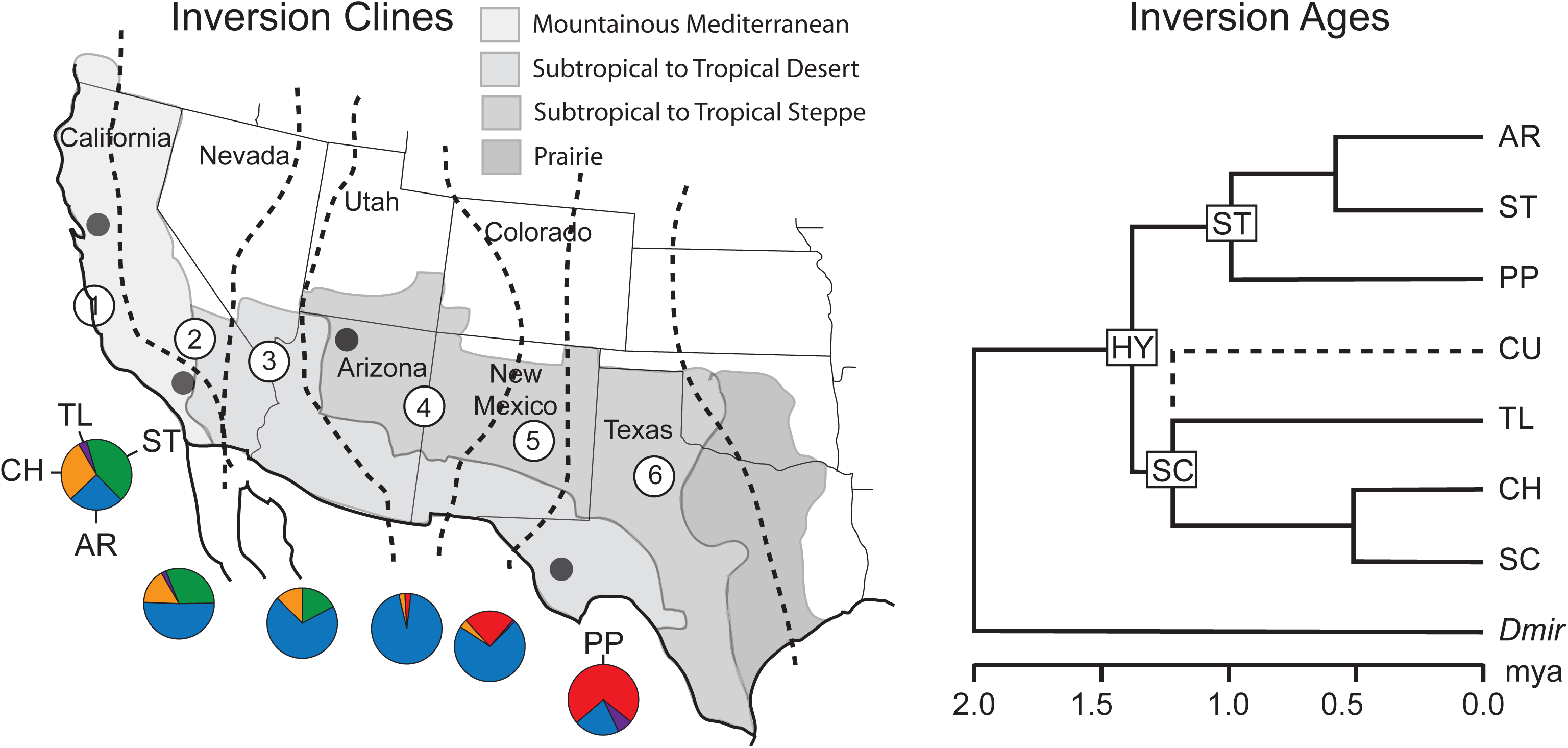
*Drosophila pseudoobscura* inversion cline and ages. The inversion abbreviations are AR, Arrowhead; CH, Chiricahua; CU, Cuernavaca; HY, Hypothetical; PP, Pikes Peak; SC, Santa Cruz; ST, Standard; and TL, Tree Line. (Left) Gene arrangement cline in *D. pseudoobscura* across the southwestern United States. The six regions delineated on the map are different climatic zones or niches inferred in Schaeffer (2008) with pie charts indicating the frequencies of the major arrangements found in each zone from 1940 inversion samples (Dobzhansky, 1944). (Right) Ages of the inversions estimated from 18 nucleotide markers sampled from the third chromosome (A. G. Wallace et al., 2011). Dmir is the outgroup *D. miranda*. The Hypothetical arrangement is the common ancestor of all arrangements within *D. pseudoobscura*. The CU arrangement is connected to the SC arrangement via a dotted line because CU is derived from SC, but was not sampled in the Wallace et al. (2011) study

The geographic clines have been stable since their initial description in the 1940s (Wyatt W. Anderson et al., 1991; Dobzhansky, 1944) despite the homogenizing effect of extensive gene flow among populations (Coyne, Bryant, & Turelli, 1987; Kovacevic & Schaeffer, 2000; Riley, Hallas, & Lewontin, 1989; Schaeffer & Miller, 1992). Altitudinal clines have also been observed (Dobzhansky, 1948a) and frequencies of the gene arrangements show seasonal cycling (Dobzhansky, 1943). In some cases, inversion frequencies in population cage experiments formed stable equilibria mimicking patterns in natural populations (Dobzhansky, 1948b, 1950; Wright & Dobzhansky, 1946). Inversion polymorphism tended to be eliminated from population cages initiated with *D. pseudoobscura* (W. W. Anderson, Dobzhansky, & Kastritsis, 1967). The problem is that karyotypic fitness can vary among different climatic zones (Schaeffer, 2008) and the Anderson et al. (1967) population cage studies do not replicate natural climates.

Allozyme and nucleotide sequence polymorphism data support the indirect effect hypothesis because different arrangements are found to have unique allelic combinations (Prakash & Lewontin, 1968, 1971; Schaeffer et al., 2003), however, these results should be viewed with caution due to the small sample sizes and limited number of loci examined. Recent transcriptomic analyses have shown that inversions capture multiple differentially expressed genes among the arrangements further supporting the indirect effect of recombination suppression hypothesis (Fuller et al., 2016).

Schaeffer (2008) used numerical analyses of a selection-migration balance model to infer the fitnesses of gene arrangement karyotypes in six niches across the east-west inversion cline (Figure 1). The fitness estimates revealed over- and under-dominance operating on the gene arrangement karyotypes in the different niches. The selection in heterogeneous environments model completely recapitulated the inversion cline from an ancestral arrangement where the frequencies of some arrangements both increased and decreased over time. The AR, PP, and CH arrangements show a steady increase to intermediate frequency from their origins suggesting that nucleotide variation at selected loci may show signatures of hard sweeps. The model also showed that ST had both increased and decreased depending on the frequency of other arrangements in the niche suggesting that patterns of ST nucleotide diversity may show a soft sweep signature. The cline observed across the southwestern United States only represents a portion of the ecological and karyotypic diversity seen in *D. pseudoobscura* with the species range extending into Mexico where the TL arrangement is found at much higher frequencies.

Inexpensive high-throughput sequencing methods now allow us to evaluate inversion establishment and maintenance hypotheses in more detail. Here, we present an analysis of 54 third chromosomes (Muller C) from six *D. pseudoobscura* gene arrangements. We used large insert mate-pair libraries to map the inversion breakpoints to test one aspect of the position effect hypothesis, i.e., whether inversion breakpoints disrupt the coding regions of genes. To test a monophyletic origin of the arrangements, phylogenetic analyses across different syntenic regions of the third chromosome were carried out. Molecular population genetic analyses tested 2,669 gene regions on the third chromosome for signatures of adaptive evolution and the structure of linkage disequilibrium was investigated across the chromosome to test for significant non-random associations as an indirect effect of suppressed recombination.

## Materials and Methods

### *Drosophila pseudoobscura* Strains

Genome sequences of 54 *D. pseudoobscura* strains were analyzed in this study. The genomes of 47 strains were described in the previous study of codon usage bias by Fuller et al. (2014). The strains were collected from the seven localities: Mount Saint Helena, CA (MSH), Santa Cruz Island, CA (SCI) (collected by Luciano Matzkin, University of Arizona), James Reserve, CA (JR) (collected by Wyatt W. Anderson, University of Georgia), Kaibab National Forest, AZ (KB), Bosque del Apache Wild Life Refuge (BdA, collected by Sara Sheeley, Upper Iowa University and provided by Bryant McAllister, University of Iowa), Davis Mountains, TX (DM), and San Pablo Etla, Oaxaca, Mexico (SPE123) (collected by Theresa A. Markow, UC San Diego). The genome sequences for seven additional strains were generated for this study. The strains carried one of six different gene arrangements: Standard (ST), Arrowhead (AR), Pikes Peak (PP), Chiricahua (CH), Cuernavaca (CU), or Tree Line (TL) (Dobzhansky & Sturtevant, 1938) (Figure 2). The strains were made homozygous for the third chromosome (Muller C, See Muller, 1940) by inbreeding, brother sister mating in a heterozygous strain, or balancer crosses (see Supporting Information).

**Figure 2.**
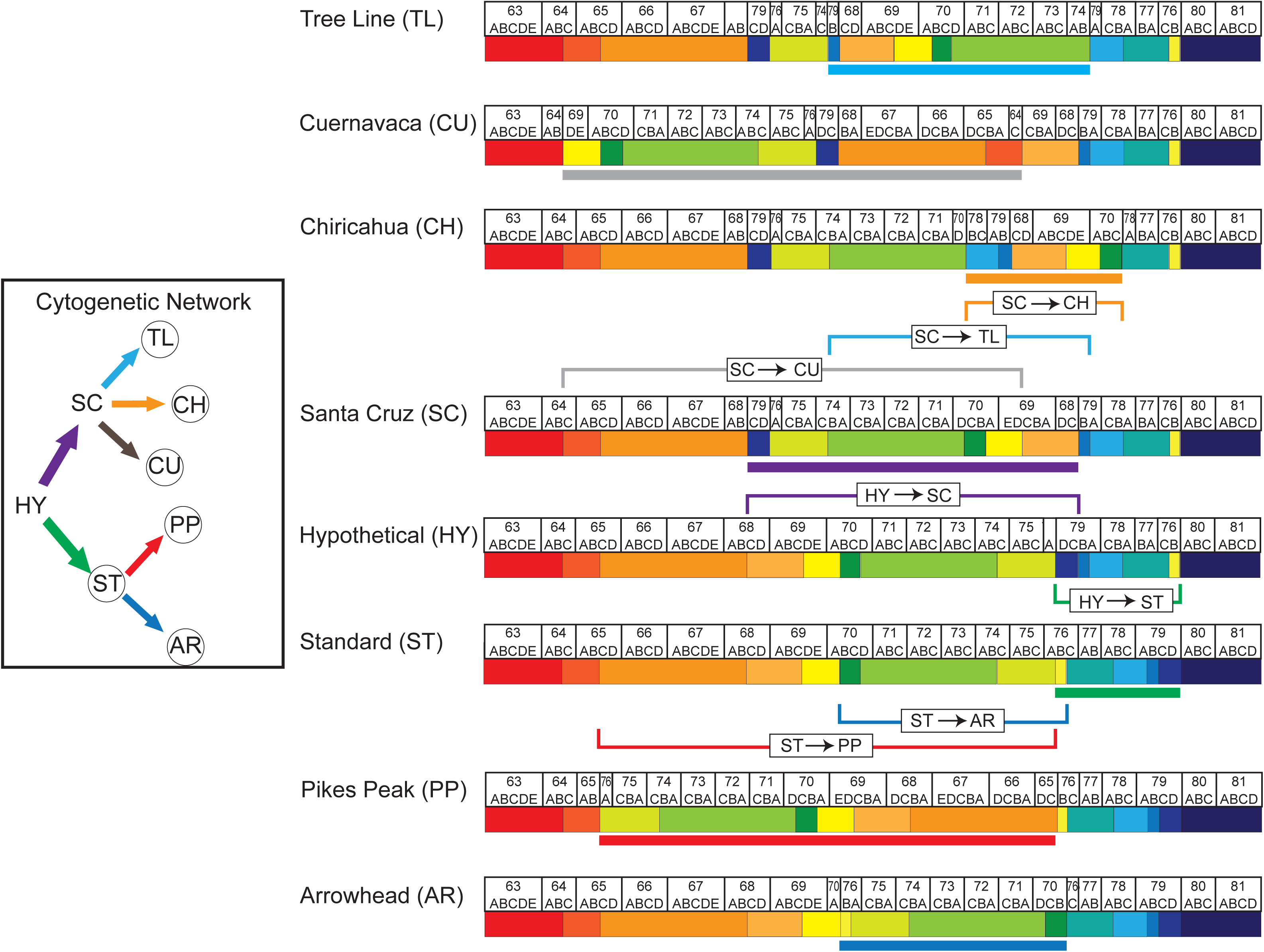
Phylogenetic network of the *Drosophila pseudoobscura* third chromosome gene arrangements inferred from polytene chromosomes isolated from larval salivary glands (Dobzhansky, 1944; Dobzhansky & Sturtevant, 1938). The box on the left shows the phylogenetic network of the different gene arrangements. The hypothetical arrangement is the ancestral arrangement (A. G. Wallace et al., 2011) and the arrows indicate the derivation of each arrangement from its ancestor. The strains with the circled arrangements were sequenced in this study. The chromosomes depicted to the right represent the numbered sections on the cytological map of *D. pseudoobscura* (Schaeffer et al., 2008). Beginning with the Hypothetical arrangement, the colored brackets indicate the segment of the ancestral arrangement that inverted and the corresponding segment is indicated with the same colored bar in the derived arrangement. These colors also match the arrows in the network in the box on the right. The colored sections under the maps indicate syntenic regions that are conserved among all gene arrangements.

### Illumina Library Construction and Sequencing

Genomic DNA samples from single male flies of each strain were purified using Qiagen DNAeasy Blood and Tissue Kit following manufacturer recommendations, including an RNAse digestion step. High molecular weight double stranded genomic DNA samples were constructed into Illumina paired end libraries according to the manufacturer’s protocol (Illumina Inc.) with modifications as described in the Supporting Information. Sequencing analysis was first done with Illumina analysis pipeline. Sequencing image files were processed to generate base calls and phred-like base quality scores and to remove low-quality reads.

### Analysis of High Throughput Sequencing Reads and Final SNP Dataset

The paired end sequence reads (101 bp) were aligned to the *D. pseudoobscura* reference strain FlyBase version 3.02 (http://flybase.org) using bwa-mem (v. 0.7.8 Li & Durbin, 2009) with default parameters. GATK (v. 3.1.1 McKenna et al., 2010) software was used to remove duplicate sequence reads, recalibrate base quality scores, and locally realign regions around indels for BWA alignments (DePristo et al., 2011). Pileup files generated with SAMtools (v. 0.1.19 Li et al., 2009) were used to determine the coverage distributions for each strain. The reference genome carries the AR arrangement (Richards et al., 2005). We called single nucleotide polymorphisms (SNPs) using the population haplotype-based software FreeBayes (v. 0.9.21 Garrison & Marth, 2012). Sites with a phred-quality score less than 30 or coverage less than two were filtered from the data. On average, 96.066 % of nucleotide sites on the third chromosome were retained for each individual strain. Because our crossing scheme produced individuals isogenic for the third chromosome, we filtered sites called as heterozygous as they are likely to be the result of read misalignment, repetitive sequence regions, or poor read mapping. Of the 19.8 Mb on the third chromosome, 0.03-0.06 % of the nucleotide sites were called as heterozygous with genotype Phred score >30. As described in Fuller et al.(2014), we found no evidence for clusters of such sites that might suggest particular regions that resisted becoming isogenic. A SNP table for the third chromosome is available through Scholarsphere (https://scholarsphere.psu.edu).

### Mapping Gene Arrangement Breakpoints

The breakpoints of the 5.9 Mb inversion that converted the Standard arrangement into Arrowhead were previously mapped using PCR (Richards et al., 2005). Large insert (3 kb) mate-pair libraries were constructed for the PP_DM1020_B, CH_JR4_L, CH_JR32_B, TL_MSH76_B, and CU_SPE123_5-2_B strains to map the locations of derived breakpoints for the inversions that converted ST to PP, HY to ST, HY to SC, SC to CH, SC to CU, and SC to TL (see Figure 2 for the schematic maps and Figure S1 in the Supporting Information for the karyotypic maps). DNA from each strain was fragmented to an average size of 3 kb using the Nextera protocol from Illumina. The 3 kb fragments were circularized and sonicated to produce 400 bp fragments. The resulting DNAs were sequenced from the 5’ and 3’ ends. The sequenced fragments included both mate pairs where the ends are separated by 3 kb and paired ends separated by 400 bp. The reads from the mate-pair library were mapped to the *D. pseudoobscura* AR reference genome (MV 2-25 version 3.02) using bwa-mem (v. 0.7.8 Li & Durbin, 2009) under default parameters. The resulting SAM file was used to locate the breakpoints (Corbett-Detig, Cardeno, & Langley, 2012). For more details about breakpoint mapping, see the Supporting Information.

### Phylogenetic Analysis of Gene Arrangements

Previous molecular evolutionary analysis of a limited number of genetic markers across the third chromosome supported a unique origin of the different gene arrangements of *D. pseudoobscura* (Aquadro et al., 1991; A. G. Wallace et al., 2011). We performed phylogenetic analysis on the more complete SNP data from across the third chromosome to test whether inversions were of unique origin. In addition, we used phylogenetic analysis of SNPs from 14 syntenic blocks across the chromosome to determine if all regions have a similar history. We used neighbor-joining as implemented in MEGA6 (Koichiro Tamura, Stecher, Peterson, Filipski, & Kumar, 2013) to infer the relationships among the 54 *D. pseudoobscura* strains using the outgroup strain, *D. miranda* (Saitou & Nei, 1987). The evolutionary distances were computed using the Maximum Composite Likelihood method (K. Tamura, Nei, & Kumar, 2004) assuming either a uniform or heterogeneous rate of evolution. All positions containing gaps and missing data were eliminated. The percentage of replicate trees in which the associated taxa clustered together were determined with 500 bootstrap replicates (Felsenstein, 1985).

### Classification of SNP sites

The inversions represent different subpopulations and we can test for genes with significant elevated frequencies of unique derived mutations, which are candidate genes targeted by adaptive evolution. The segregating sites discovered in this sample are a composite of unique and shared mutations. A unique derived mutation was one that occurred on the branch immediately before or after the inversion mutation. Shared mutations either predate two or more inversion mutations or are alleles transferred among arrangements via gene conversion or double cross overs (Arcadio Navarro, Betrán, Barbadilla, & Ruiz, 1997). Shared polymorphisms could also result from the balancer crosses that were used to generate the isochromosomal strains (Miller et al., 2016). See the Supporting Information for more details on how polymorphic sites were classified.

### Estimates of Site Frequency Spectrum and Mean Derived Allele Frequency within Protein Coding Genes within Chromosomal Arrangements

A total of 2,669 protein coding gene models are annotated for the third chromosome in release Dpse 3.02 FB14_04 in FlyBase (http://flybase.org). We examined the site frequency spectrum and mean derived allele frequency (DAF) for each gene including 1,000 bps upstream and downstream of the annotated transcriptional start and end sites. We used the largest of multiple encoded transcripts for each gene. The site frequency spectrum for each gene was summarized with the ratio of Tajima’s *D* (1989) to the absolute value of its theoretical minimum (*D*_min_), which occurs when all segregating sites have a frequency of 1/*n*, where *n* is the sample size. Tajima’s *D* is sensitive to sample size and numbers of segregating sites, but *D*/|*D*_min_| controls for arrangements with different sample sizes (Schaeffer, 2002). The *D*/|*D*_min_| has a minimum value of −1 and its mean value will reflect the demographic history of the population, which should be similar for all loci in the genome (Hahn, Rausher, & Cunningham, 2002; Schaeffer, 2002). We used a random permutation test to detect outlier loci that have clusters of low or high frequency variants. Shuffling the position of sites will break up these clusters such that the observed values of *D/*|*D*_min_| will be more extreme than the values based on random permutations for a given number of segregating sites and the mean local site frequency spectrum in proximal, inverted, or distal regions. The segregating sites within each region were shuffled without replacement 100,000 times and we estimated how often the permuted Tajima’s *D*/|*D*_min_| was more extreme than the observed value in the gene region. This approach maintained the linkage relationships of variable nucleotides within the chromosome region. We used the *q*-value method with a false discovery rate (FDR) of 0.01 to correct for multiple tests (Storey, 2002, 2003). Shared and unique SNPs were analyzed separately for the five gene arrangements with sample sizes greater than eight, Arrowhead (15), Standard (8), Pikes Peak (10), Chiricahua (9), and Tree Line (9). A similar approach was used to detect significant clusters of high frequency derived unique mutations per segregating site (DAF) in the 2,669 genes, which includes polymorphic and fixed mutations.

### Population Specific Branch Length Tests

A second method to detect outlier genes that accumulated large numbers of arrangement specific mutations is similar to other statistics such as the “Population Branch Statistic” (PBS; Yi et al., 2010) and “Locus Specific Branch Length” (LSBL; Shriver et al., 2004). This test has been used to identify loci with evidence of adaptive evolution (Huerta-Sanchez et al., 2013). Prior use of PBS and LSBL require exactly three subpopulations. Here, we extend these methods to an unrooted phylogeny of known topology containing any number of branches and nodes and details of the analysis can be found in Supporting Information.

We estimated the Population Specific Branch Length (PSBL) for each gene on the third chromosome including 1 kb upstream and downstream. We used similar bootstrap approaches as our analyses of *D*/|*D*_min_| and DAF, to detect significantly large PSBL estimates with a false discovery rate (FDR) of 0.01 (Storey, 2002, 2003).

The detection of statistical outliers could be performed with coalescent simulations which specify the demographic history of each arrangement. We know the branching order of the different gene arrangements, but we have insufficient information about the demography of each arrangement to carefully specify a precise model for the system of inversions. For this reason, we used random permutation tests that shuffled SNP positions and looked for genes that were statistical outliers rather than searching for parameter values that could support any particular model or hypothesis.

### Detection of Putative Functional Variation in Protein Coding Genes Within Gene Arrangements

We extracted and translated the 2,669 third chromosome coding sequences from the *D. pseudoobscura* reference annotation version 3.02 available from FlyBase, (dos Santos et al., 2015). We used the same scoring scheme that was used to classify nucleotide polymorphisms (Figure S2 in the Supporting Information). Variation in stop codons was counted in the data set. We inferred the amino acid transition matrix counting the number of events from the ancestral to derived amino acids. Additionally, we examined the frequency spectrum of the amino acid transitions for all sites.

### Detection of Significant Linkage Disequilibrium

Linkage disequilibrium was characterized for all polymorphic sites across the third chromosome using the correlation based approach of Zaykin *et al.* (2008), similar to *r*^2^ (Hill & Robertson, 1968). A heat map image matrix was generated by taking the average significance values at 100 adjacent sites. At each polymorphic site, we also performed a Fisher’s exact test to assess the significance of allele association within chromosomal arrangements. All significance values were corrected for multiple testing by controlling the FDR (Storey, 2003).

## Results

### Map of Inversion Breakpoints

Thirty eight genetic and physical markers from Muller C were used to bracket the locations of each breakpoint in the AR reference sequence (see Supplemental Table 21 Muller C tab, Schaeffer et al., 2008). Seven pairs of inversion breakpoints were mapped with the mate pair data similar to previous approaches (Corbett-Detig et al., 2012; Cridland & Thornton, 2010)(See Supporting Information). Mate pair reads that mapped at megabase distances apart are candidates for breakpoint positions if the two ends coincide with the approximate locations of breakpoints on the cytological map (Dobzhansky, 1944; Dobzhansky & Sturtevant, 1938).

All breakpoints mapped within intergenic regions and not within the transcripts of the boundary genes (Table 1). An average of ∼4.5 breaks (95% CI: 1-8) are expected in intergenic regions if 14 breakpoints are placed randomly, using a uniform distribution, on the third chromosome map that includes 6.3 Mb of intergenic regions and 13.4 Mb of coding nucleotides. Four of the thirteen breakpoint regions had nucleotide sites with elevated coverage (pHYSC, pSTAR, dSTAR, and pSCCH). We failed to observe elevated read coverage in any boundary gene indicating that genes adjacent to inversion breakpoints are not duplicated as has been observed in other Drosophila breakpoints (Calvete, Gonzalez, Betran, & Ruiz, 2012; Guillen & Ruiz, 2012; Papaceit, Segarra, & Aguadé, 2013; Puerma et al., 2014; Ranz et al., 2007).

**Table 1.** Breakpoint map locations for seven inversion events in the MV 2-25 reference genome of *D. pseudoobscura.*

| Breakpoint | Cyto_Ref | 5'_Gene | 5'_Coor | 3'_Gene | 3'_Coor | BP_Interval |
| --- | --- | --- | --- | --- | --- | --- |
| pSCCU | 64B\|64C | GA17643 | 1,766,685 | GA24634 | 1,769,869 | 1,769,503..1769,545 |
| pSTPP | 65B\|65C | GA13487 | 2,500,7218 | GA13475 | 2,505,628 | 2,504,717..2,505,525 |
| pHYSC | 68B\|68C | GA21475 | 6,814,856 | GA19795 | 6,830,821 | 6,817,305..6,819,934 |
| dSCCU | 69C\|69D | GA18750 | 8,021,189 | GA24813 | 8,027,622 | 8,026,967..8,027,480 |
| pSTAR | 70A\|76B | GA24334 | 8,902,967 | GA19155 | 8,905,717 | 8,902,967..8,905,362 |
| dSTPP | 76B\|76A | GA22082 | 9,162,365 | GA20777 | 9,164,720 | 9,163,072..9,164,670 |
| pHYST | 76B\|76A | GA22082 | 9,162,375 | GA20777 | 9,164,720 | 9,163,072..9,164,057 |
| pSCTL | 74C\|74B | GA19582 | 10,623,204 | GA14679 | 10,628,103 | 10,626,931..10,627,448 |
| pSCCH | 70D\|70C | GA20939 | 13,940,052 | GA12507 | 13,941,493 | 13,940,125..13,941,042 |
| dSTAR | 70B\|76C | GA17716 | 14,813,423 | GA17559 | 14,833,672 | 14,831,322..14,833,071 |
| dSCCH | 78A\|78B | GA10355 | 15,653,239 | GA13882 | 15,658,703 | 15,657,098..15,657,203 |
| dSCTL | 79B | GA13539 | 17,149,432 | GA24132 | 17,153,461 | 17,151,554..17152,967 |
| dHYSC | 79B\|79C | GA21720 | 17,545,780 | GA24111 | 17,567,890 | 17,545,438..17,545,671 |
| dHYST | 79D\|80A | GA30270 | 17,725,786 | GA10842 | 17,729,964 | 17,727,100..17,727,669 |

Breakpoint, is the first letter indicates if the breakpoint is proximal (p) or distal (d), the second two letters indicates the ancestral gene arrangement and the last two letters indicates the derived gene arrangement. Cyto, is the location of the breakpoint in the AR reference genome. The location of the dSTPP breakpoints differ from the original location mapped in Dobzhansky and Sturtevant (1938). The original map location of dSTPP was 75C|76A. 5'\_Gene, the FlyBase annotation symbol of the gene 5' to the breakpoint. 5'\_Coor, the nucleotide coordinate of the last base of the 5'\_Gene. 3'\_Gene, the FlyBase annotation symbol of the gene 3' to the breakpoint. 3'\_Coor, the nucleotide coordinate of the first base of the 3'\_Gene. BP\_Interval, the coordinates of the breakpoint region in the AR reference sequence defined by mate pair reads spanning the inversion breakpoint. All coordinates are those of the AR reference genome (MV 2-25).

**Table 2.**
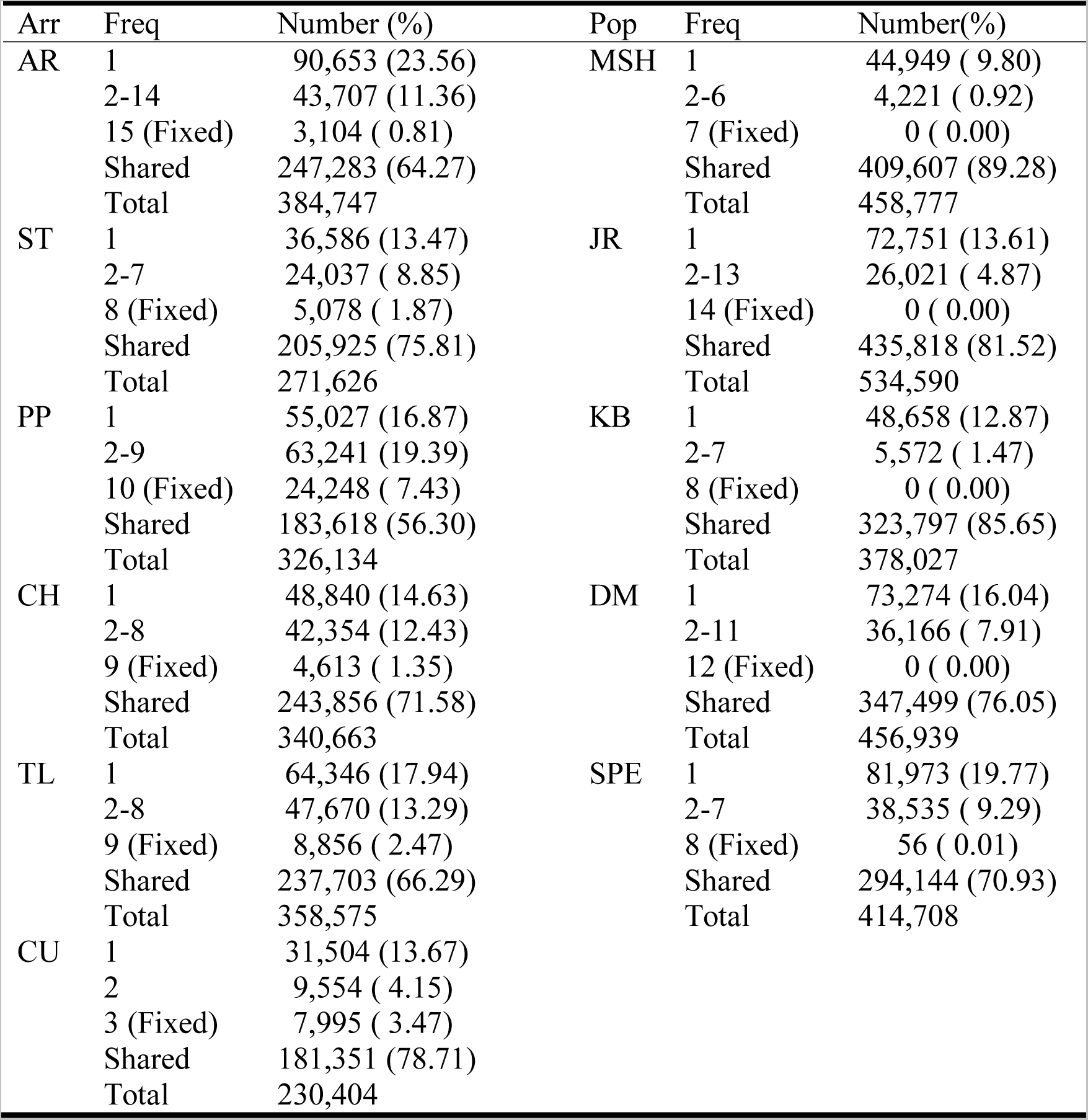
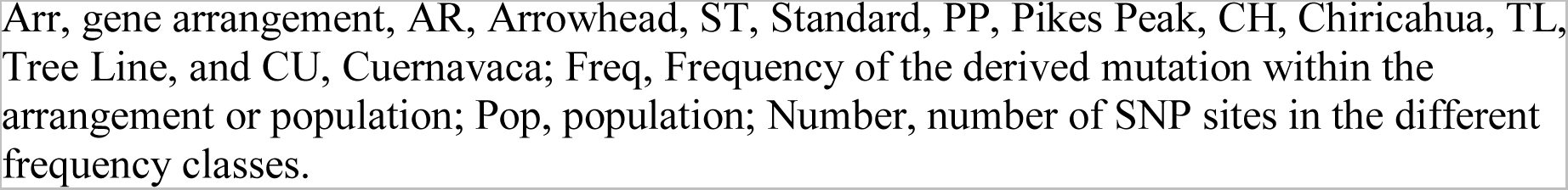
Site frequency spectrum analysis based on gene arrangement and population categorical variables.

| Arr | Freq | Number (%) | Pop | Freq | Number(%) |
| --- | --- | --- | --- | --- | --- |
| AR | 1 | 90,653 (23.56) | MSH | 1 | 44,949 ( 9.80) |
|  | 2-14 | 43,707 (11.36) |  | 2-6 | 4,221 ( 0.92) |
|  | 15 (Fixed) | 3,104 ( 0.81) |  | 7 (Fixed) | 0 ( 0.00) |
|  | Shared | 247,283 (64.27) |  | Shared | 409,607 (89.28) |
|  | Total | 384,747 |  | Total | 458,777 |
| ST | 1 | 36,586 (13.47) | JR | 1 | 72,751 (13.61) |
|  | 2-7 | 24,037 ( 8.85) |  | 2-13 | 26,021 ( 4.87) |
|  | 8 (Fixed) | 5,078 ( 1.87) |  | 14 (Fixed) | 0 ( 0.00) |
|  | Shared | 205,925 (75.81) |  | Shared | 435,818 (81.52) |
|  | Total | 271,626 |  | Total | 534,590 |
| PP | 1 | 55,027 (16.87) | KB | 1 | 48,658 (12.87) |
|  | 2-9 | 63,241 (19.39) |  | 2-7 | 5,572 ( 1.47) |
|  | 10 (Fixed) | 24,248 ( 7.43) |  | 8 (Fixed) | 0 ( 0.00) |
|  | Shared | 183,618 (56.30) |  | Shared | 323,797 (85.65) |
|  | Total | 326,134 |  | Total | 378,027 |
| CH | 1 | 48,840 (14.63) | DM | 1 | 73,274 (16.04) |
|  | 2-8 | 42,354 (12.43) |  | 2-11 | 36,166 ( 7.91) |
|  | 9 (Fixed) | 4,613 ( 1.35) |  | 12 (Fixed) | 0 ( 0.00) |
|  | Shared | 243,856 (71.58) |  | Shared | 347,499 (76.05) |
|  | Total | 340,663 |  | Total | 456,939 |
| TL | 1 | 64,346 (17.94) | SPE | 1 | 81,973 (19.77) |
|  | 2-8 | 47,670 (13.29) |  | 2-7 | 38,535 ( 9.29) |
|  | 9 (Fixed) | 8,856 ( 2.47) |  | 8 (Fixed) | 56 ( 0.01) |
|  | Shared | 237,703 (66.29) |  | Shared | 294,144 (70.93) |
|  | Total | 358,575 |  | Total | 414,708 |
| CU | 1 | 31,504 (13.67) |  |  |  |
|  | 2 | 9,554 ( 4.15) |  |  |  |
|  | 3 (Fixed) | 7,995 ( 3.47) |  |  |  |
|  | Shared | 181,351 (78.71) |  |  |  |
|  | Total | 230,404 |  |  |  |

Arr, gene arrangement, AR, Arrowhead, ST, Standard, PP, Pikes Peak, CH, Chiricahua, TL, Tree Line, and CU, Cuernavaca; Freq, Frequency of the derived mutation within the arrangement or population; Pop, population; Number, number of SNP sites in the different frequency classes.

### Phylogenetic Relationships of Arrangements 14 Syntenic Regions Show Diverse Histories across the Third Chromosome

We used paired-end reads generated from each strain to identify single nucleotide polymorphisms (SNPs). The median coverage across all strains was 35 (Table S4 and Figure S24). From these aligned reads, we identified 1,028,037 SNPs on the third chromosome and estimated error rates to vary from 9.7 x 10^-4^ in AR to 3.4 x 10^-3^ in PP, CH, and TL using a set of previously sequenced markers (Tables S5 and S6).

The overall phylogenetic tree supports the unique origin hypothesis based on the monophyletic relationship of each of the six arrangements (Figure 3) consistent with RFLP and short nucleotide marker data (Aquadro et al., 1991; A. G. Wallace et al., 2011). The root of the tree, HY, was also confirmed with gene adjacency information at the breakpoints (Bhutkar et al., 2008)(See Identification of Inversion Breakpoints in Supporting Information). The branching order is consistent with the cytogenetic phylogeny, except that PP is inferred to be the sister clade to the AR, ST, CH, CU, and TL gene arrangements rather than forming a monophyletic group with ST and AR (Dobzhansky & Sturtevant, 1938).

**Figure 3.**
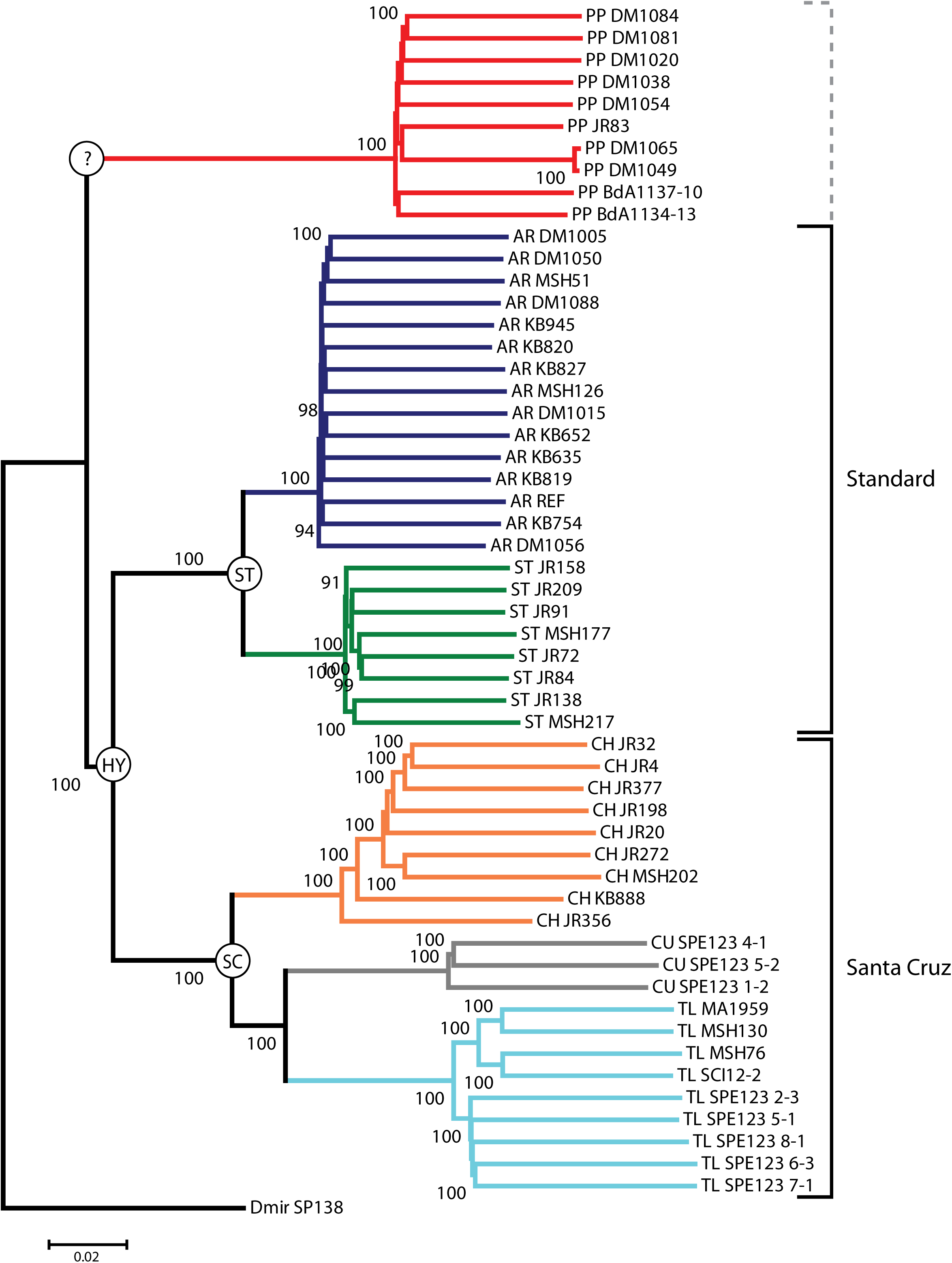
Phylogeny inferred with the neighbor-joining method (Saitou & Nei, 1987). A total of 1,028,037 SNPs were used in each of the regions to construct the gene arrangement phylogeny. A total of 500 bootstrap replicated were used to determine the confidence in the nodes of the trees where only nodes with > 90% support shown on the tree.

We tested whether PP showed this unexpected relationship in all regions of the third chromosome (Figure 2). We used neighbor-joining to infer the relationships among the six arrangements in the 14 syntenic regions (Figure 4, Figures S25-S38 in the Supporting Information). The six gene arrangements form monophyletic clusters in the central syntenic blocks where recombination is reduced. Not all syntenic blocks, however, are concordant with the established cytogenetic phylogeny, e.g., see regions 68C-69C, 69D-70A, 76C-78A, and 78B-79A in Figure 4 compared with the cytogenetic phylogeny in (Figure 2). PP consistently diverges before the split of the AR and ST lineages within the ST phylad in these four regions. The branching order of the CH, CU, and TL arrangements within the Santa Cruz phylad is also not consistent among these four syntenic regions. While CU is always the most derived member of the Santa Cruz phylad, it is not clear whether CH or TL is the more ancestral member.

**Figure 4.**
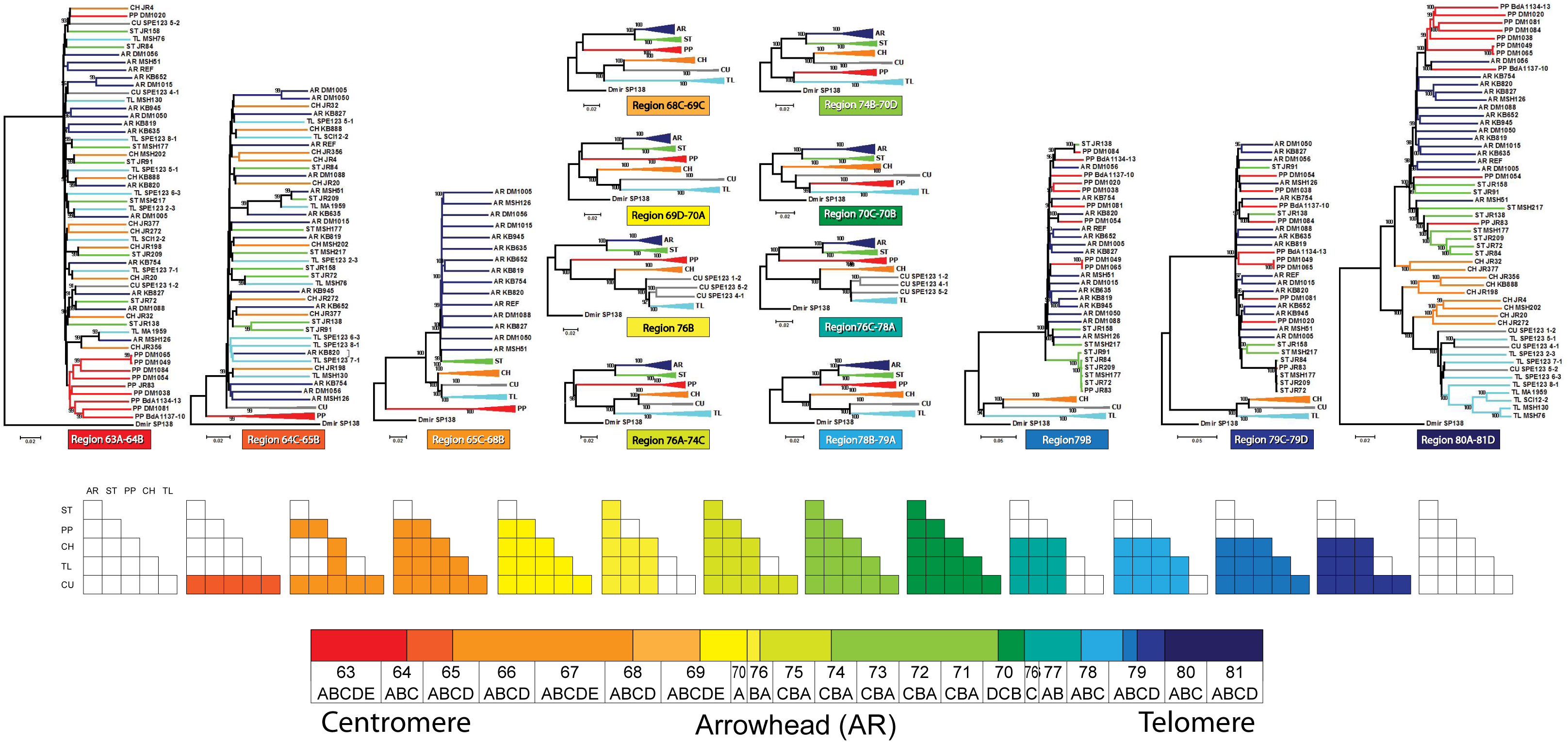
Top Row: Phylogenies inferred with the neighbor-joining method (Saitou & Nei, 1987). Phylogenies were inferred in each of 14 syntenic blocks defined by seven pairs of inversion breakpoints. Below each phylogeny is a colored block with a label that corresponds to the 14 syntenic blocks on the cytogenetic map shown at the bottom. Indels were removed from the analysis. The following numbers of SNPs were used in each of the regions to construct the phylogenies: 63A-64B, 36,372 SNPs; 64C-65B, 24,361 SNPs; 65C-68B, 239,432 SNPs; 68C-69C, 77,386 SNPs; 69D-70A, 57,145 SNPs; 76B, 16,556 SNPs; 76A-74C, 91893 SNPs; 74B-70D, 203,463 SNPs; 70C-70B, 59,084 SNPs; 76C-78A, 51,476 SNPs; 78B-79A, 75,395 SNPs; 79B, 18,542 SNPs; 79C-79D, 9,324 SNPs; and 80A-81D, 67,608 SNPs. A total of 500 bootstrap replicated were used to determine the confidence in the nodes of the trees where only nodes with > 90% support shown on the tree. Middle Row: The 14 matrices indicate whether a particular region is outside (open box) or within (filled box) the inverted region in a heterozygote for a particular pair of arrangements (shown on the horizontal and vertical axis). The fill colors match the colors for the syntenic blocks of the reference sequence. Bottom Row: A diagram of the cytogenetic map of the Arrowhead reference genome with colored segments representing the 14 syntenic blocks.

In syntenic regions that are discordant with the cytogenetic phylogeny, the PP clade is the most inconsistent group. PP clusters with the Santa Cruz phylad in four regions (76B, 76A-74C, 74B-70D, 70C-70B) and is basal to the phylad in two of the four regions (76B, 76A-74C). PP is sister to TL in the other two blocks (74B-70D and 70C-70B). Block 74B-70D is also unusual because the CH and CU clades cluster with the Standard phylad. In the three remaining syntenic blocks (79B, 79C-79D, 80A-81D), PP strains fail to form monophyletic groups and clusters among the AR and ST strains.

### The Site Frequency Spectrum of Inversion-Specific Mutations

We examined the site frequency spectra of inversion-specific mutations with Tajima’s *D*/|*D*_min_| (Figure 5). The frequency spectra shows more high frequency variants near and within inverted regions and an excess of rare variants in the proximal and distal regions. Proximal and distal regions have more shared than unique polymorphisms and the frequency of derived mutations is higher in shared versus unique polymorphisms with random permutation tests in five arrangements (All arrangements and regions *P* < 1.0 x 10^-3^)(Table 3) and Chi square tests of homogeneity (AR, X^2^=12,688.9, df=2, *P*< 1x10^-6^; PP, X^2^=14,562.0, df=2, *P*< 1x10^-6^; CH, X^2^=8,923.5, df=2, *P*< 1x10^-6^; TL, X^2^=12,495.1, df=2, *P*< 1x10^-6^; CU, X^2^=457.8, df=2, *P*< 1x10^-6^). ST has an excess of unique SNPs in the proximal region compared to the inverted and distal regions (ST, X^2^=1,345.1, df=2, *P*< 1x10^-6^).

**Table 3.** Unique and Shared polymorphisms in proximal, inverted, and distal regions in six gene arrangements.

| Arrangement | Type | Proximal<br>Obs (Exp) | Inverted<br>Obs (Exp) | Distal<br>Obs (Exp) |
| --- | --- | --- | --- | --- |
| Arrowhead* | Shared | 114,700 (104,926.2) | 52,305 ( 67,343.1) | 58,155 (52,890.6) |
|  | Unique | 53,908 ( 63,681.8) | 55,910 ( 40,871.9) | 26,836 (32,100.4) |
| Standard* | Shared | 141,750 (143,519.3) | 28,261 ( 28,761.2) | 18,563 (16,293.5) |
|  | Unique | 51,594 ( 49,824.7) | 10,485 ( 9,984.8) | 3,387 ( 5,656.5) |
| Pikes Peak* | Shared | 14,064 ( 11,154.7) | 96,036 (111,515.7) | 53,828 (41,257.5) |
|  | Unique | 6,696 ( 9,605.3) | 111,505 ( 96,025.3) | 22,956 (35,526.5) |
| Chiricahua* | Shared | 141,777 (131,368.2) | 49,533 ( 60,142.2) | 31,078 (30,877.6) |
|  | Unique | 46,509 ( 56,917.8) | 36,667 ( 26,057.8) | 13,178 (13,378.4) |
| Tree Line* | Shared | 104,174 ( 92,840.4) | 63,446 ( 78,297.3) | 46,432 (42,914.3) |
|  | Unique | 40,719 ( 52,052.6) | 58,750 ( 43,898.7) | 20,543 (24,060.7) |
| Cuernavaca* | Shared | 6,360 ( 5,626.2) | 105,495 (106,453.1) | 51,849 (51,624.6) |
|  | Unique | 947 ( 1,680.8) | 32,760 ( 31,801.9) | 15,198 (15,422.4) |

\*, $P < 0.0001$ in a Chi Square test of homogeneity.

**Figure 5.**
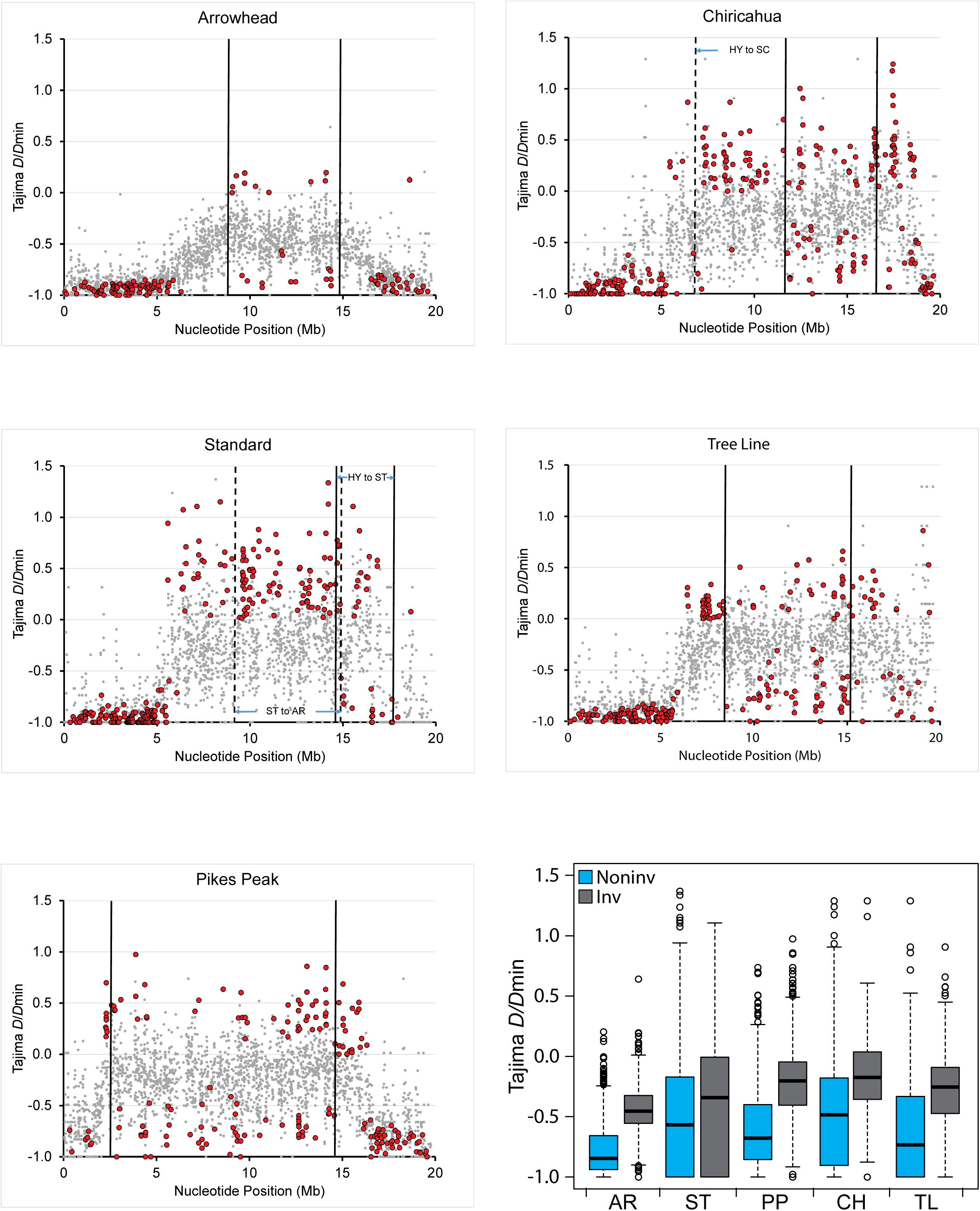
Estimates of Tajima’s *D*/|*D*_min_| estimated from unique mutations in 2,669 gene regions across the third chromosome (Muller C) of *D. pseudoobscura* in five gene arrangements, Arrowhead, Standard, Pikes Peak, Chiricahua, and Tree Line. Outlier gene regions that showed a significant negative or positive value of Tajima’s *D*/|*D*_min_| are indicated with a red point. Gene regions that did not have an extreme value of Tajima’s *D*/|*D*_min_| are shown with a gray dot. The order of genes is specific to the particular gene arrangement background. The locations of inversion breakpoints are shown with a dotted line. Box plots for all *D*/|*D*_min_| values across the third chromosome are shown in the lower right hand corner for each of the arrangements (Hintze & Nelson, 1998).

We tested for outlier genes that either have significant excesses of low (*D*/|*D*_min_|<0) or intermediate (*D*/|*D*_min_|>0) frequency SNPs. The majority of extremely low values of *D*/|*D*_min_| are found in the proximal and distal regions where shared mutations with higher frequencies were removed leaving younger low frequency mutations. Significant elevations in *D*/|*D*_min_| occurred for genes within the inversion where fewer shared polymorphisms were removed. For ST and CH, the elevation of Tajima’s *D/*|*D*_min_| extends approximately five and seven Mb upstream of the proximal breakpoint, respectively.

We estimated *D*/|*D*_min_| for the non-inverted second chromosome (McGaugh et al., 2012). Tajima’s *D/*|*D*_min_| varies uniformly across chromosome two with a mean *D*/|*D*_min_| of −0.55, which is consistent with a population expansion parameter (*Nr*) of 30 based on coalescent simulations (McGaugh et al., 2012; Schaeffer, 2002) (See Supporting Information and Figure S39). Fifteen regions show a significantly excess of intermediate frequency variants and thirty regions have a significant excess of rare variants (Figure S39). This indicates the pattern of Tajima’s *D/*|*D*_min_| across the inverted third chromosome is influenced by the presence of inversions.

The mean values of Tajima’s *D*/|*D*_min_| for the five arrangements are AR =−0.72, ST = −0.53, PP= −0.33, CH= −0.41, and TL = −0.49. The *D/*|*D*_min_| values for ST, CH, and TL are similar to that of the second chromosome consistent with a similar demographic history. The distribution of *D/*|*D*_min_| for AR is more negative than the other arrangements suggesting a greater rate of expansion for this chromosome.

### Detection of Protein Coding Regions with Elevated Frequencies of Derived Mutations

The site frequency spectrum as summarized by Tajima’s *D*|*D*_min_| does not include nucleotide sites that are fixed within a gene arrangement background in our sample. We estimated the mean derived allele frequencies (DAF) of unique polymorphisms in the 2,669 genes and tested loci for significant clusters of high frequency alleles using random permutation tests (Figure 6). The AR, ST, PP, CH, and TL arrangements had 138, 229, 233, 161, and 173 genes, respectively, with a significant elevation of mean DAF per segregating site. The mean DAF tends to be at its lowest near the centromere and telomere and increases near the inversion breakpoints with the highest values observed within the inverted regions and up to 1 Mb upstream or downstream. The exceptions are ST and CH where the mean DAF is elevated within five to seven Mb of the proximal breakpoint, which overlaps with the inverted region of AR. The mean DAF frequency is uniformly low across chromosome two (∼0.2) not reaching the high values seen for any arrangement on the third chromosome (See Figure S39 in Supporting Information).

**Figure 6.**
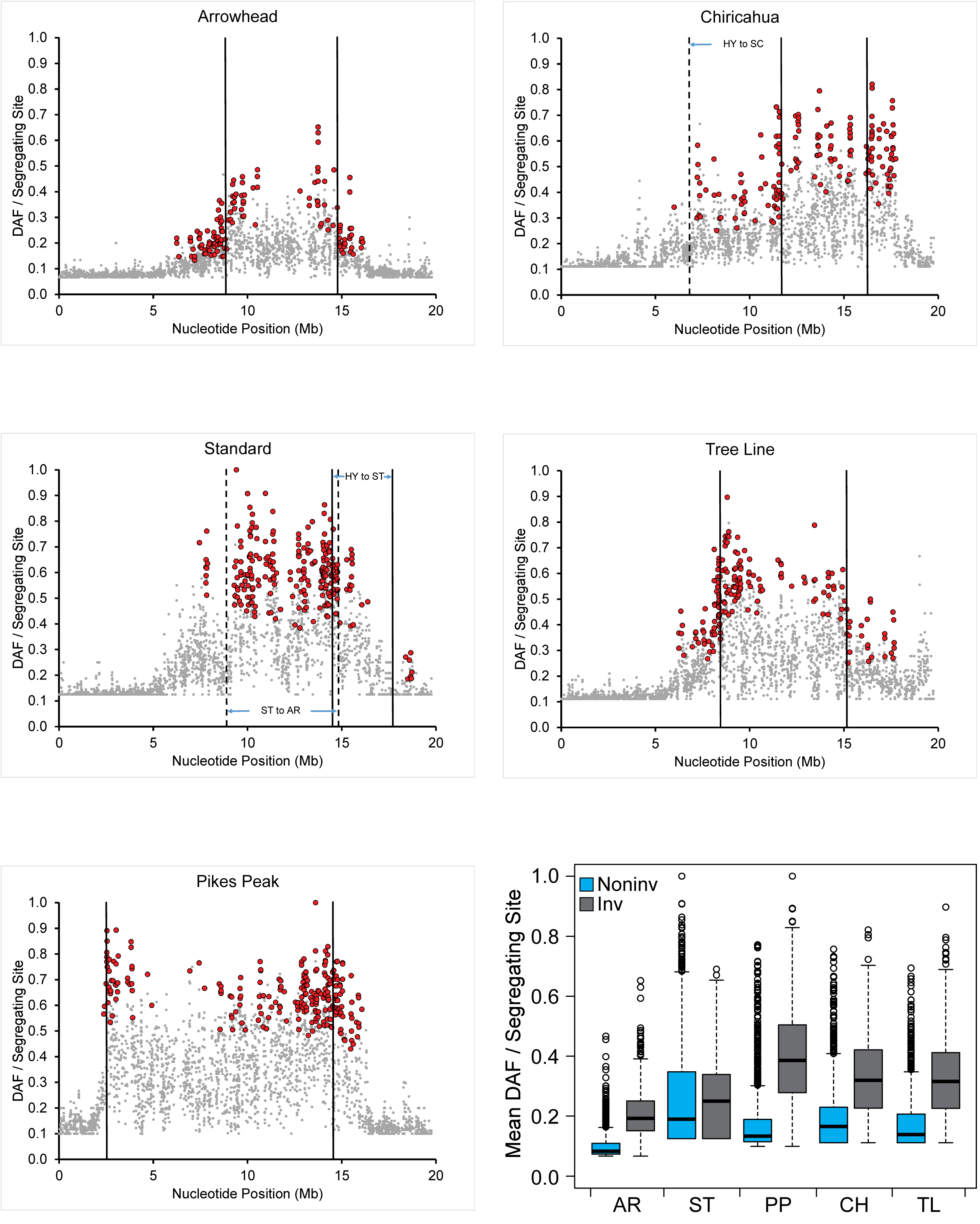
Estimates of the mean derived allele frequency (DAF) per segregating site estimated from unique mutations in the five gene arrangements, Arrowhead, Standard, Pikes Peak, Chiricahua, and Tree Line. Regions that showed a significantly large mean given the number of SNPs are indicated with a red point. Gene regions that did not have an extreme mean DAF value are shown with a gray dot. The locations of inversion breakpoints are shown with a solid or dotted line. The order of genes is specific to the particular gene arrangement background. Box plots for all mean DAF values across the third chromosome are shown in the lower right hand corner for each of the arrangements (Hintze & Nelson, 1998).

### Detection of Arrangement Specific Allele Frequency Changes with Population Specific Branch Length Analysis

Population specific branch length (PSBL) analysis allowed us to determine which gene regions have a significantly high proportion of arrangement specific allele frequency changes (see Figure 7 and Supplemental Spreadsheet 1). A significantly long branch will occur if a gene has an excess of arrangement specific mutations relative to the other four arrangements. For each lineage on the phylogeny, intervals with the longest branch lengths are located within or near inversion breakpoints. The mean branch length for genes in inverted regions is significantly greater than for genes located outside of the breakpoints in every case (Wilcoxon rank-sum test, *P*<0.05). After correcting for multiple testing with a FDR of 0.01, we determined that 317, 380, 396, 350, and 340 genes had a significantly long branch length in the AR, ST, PP, and CH arrangements, respectively. Of these, 71, 130, 192, 74, and 75 genes had long branches only within AR, ST, PP, CH, and TL, respectively. Some genes had longer branches in multiple arrangements with 249, 149, 54, and 16 genes in two, three, four, or five arrangements, respectively. A total of 1,659 genes did not have a significantly long branch length in any of the five arrangements, while 1,010 genes had a significantly long branch length in at least one arrangement.

**Figure 7.**
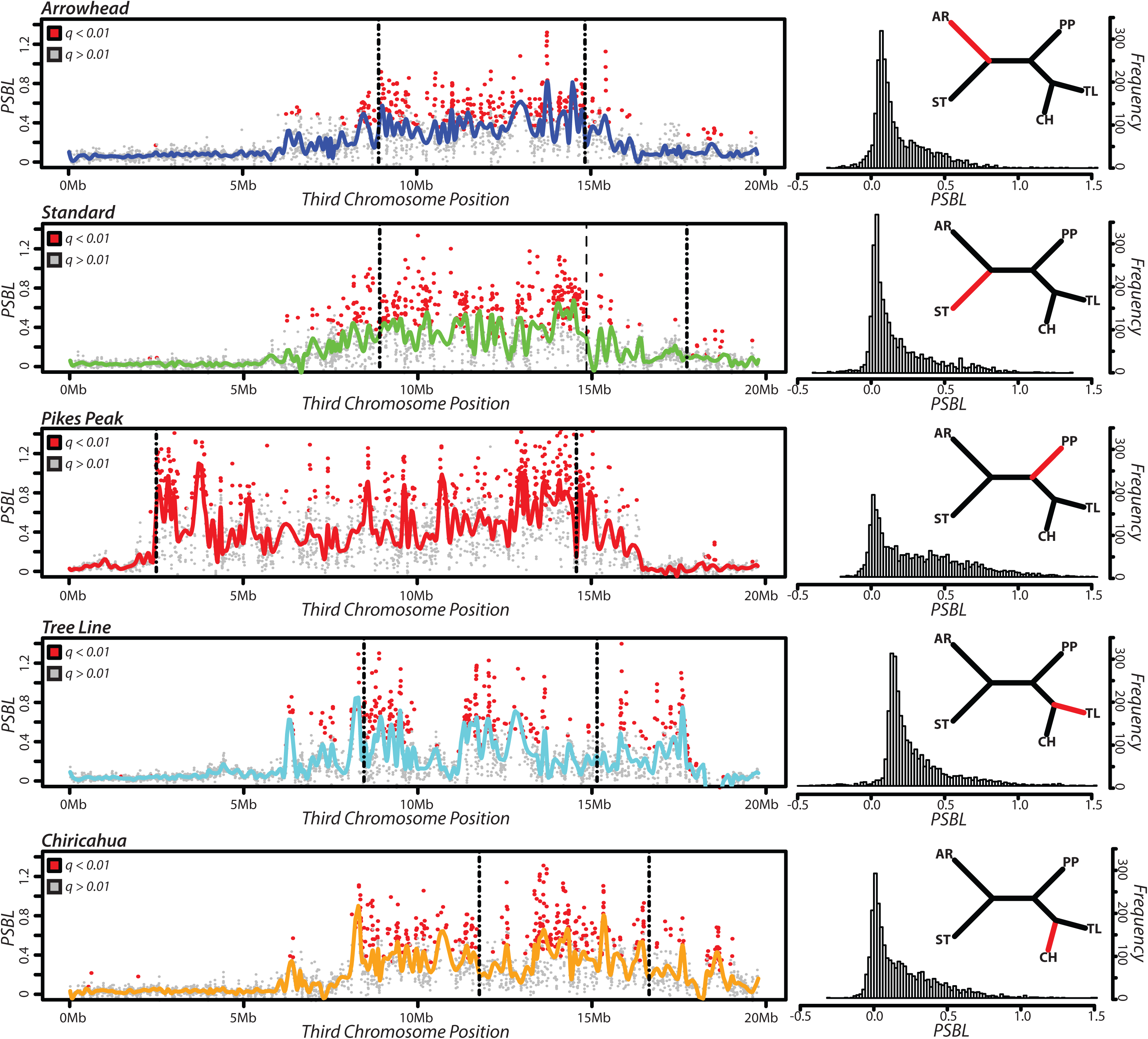
Population specific branch lengths estimated for five of the six arrangements. The plots to the left show the estimates of PSBL for the 2,669 genes across the third chromosome. Each gene is represented by a point, with those colored in red classified as statistical outliers after correcting for multiple testing. The histograms to the right show the distribution of PSBL for each arrangement with the branch highlighted in red.

### Elevated Derived Allele Frequencies and Fixed Amino Acid Changes within Inverted Gene Regions

Here, we ask how many of the DAF and PSBL outlier gene regions also harbor at least one fixed amino acid change within a particular arrangement in our sample. A total of 277 gene regions DAF and PSBL outlier genes contained at least one fixed amino acid difference in at least one gene arrangement. There are 28, 74, 144, 31, and 47 outlier genes with at least one fixed amino acid in AR, ST, PP, CH, or TL, respectively. Chi-square tests of homogeneity support a heterogeneous distribution of candidate selected genes with more outliers in inverted regions for all arrangements except ST (AR, X^2^=24.1, df=2, *P*=5.9x10^-6^; ST, X^2^=9.2, df=2, *P*=0.010; PP, X^2^=36.1, df=2, *P*=1.4x10^-8^; CH, X^2^=28.7, df=2, *P*=5.7x10^-7^; TL, X^2^=42.2, df=2, *P*=6.9x10^-10^).

The outlier genes are found across the length of the inverted regions, but are not uniformly distributed based on Kolmogorov-Smirnov tests (AR *D*_max_=-0.258 P=0.005; ST, *D*_max_=0.387 P=0.001; PP *D*_max_=0.387 P<1 x 10^-4^; CH *D*_max_=-0.285 P=0.001; TL *D*_max_=-0.555 P<1 x 10^-4^).The distance of the closest outlier gene to the center of the inverted region in AR, ST, PP, CH, and TL is 0.92, 0.62, 0.04, 0.08, and 0.11 Mb, or 15.5, 20.0, 0.3, 1.7, and 1.7 % of the total inversion size, respectively. Genes with the maximum DAF frequency or PSBL are more centrally located within the inverted region and are not adjacent to either the proximal or distal breakpoints in each case.

We tested for a linear association between the size of each chromosomal region (proximal, inverted, and distal) and the proportion of genes showing evidence of adaptive evolution (Table 4), observing a significant positive relationship (*P*=0.003, *R*^2^=0.96). While there is a positive correlation in proximal and distal regions, the relationship is not significant (proximal: *P*=0.14, *R*^2^=0.57; distal: *P*=0.37, *R*^2^=0.27). In four gene arrangements (AR, PP, TL, CH), there is an excess of candidate adaptive genes within inverted regions.

**Table 4.** Relationship between length of three chromosomal regions and the percent of selected protein coding genes within the regions.

| Region | N | Intercept (SE) | Slope (SE) | *F* | *P* | *R*<sup>2</sup> |
| --- | --- | --- | --- | --- | --- | --- |
| Proximal | 5 | -1.08 (1.11) | 0.22 (0.11) | 4.00 | 0.14 | 0.57 |
| Inverted | 5 | 0.77 (0.79) | 1.55 (0.19) | 66.56 | 0.003 | 0.96 |
| Distal | 5 | 3.48 (0.77) | 0.53 (0.50) | 1.11 | 0.36 | 0.27 |

We asked if recombination rates varies within and outside of outlier gene regions using the fine-scale maps of the population scaled recombination rate ρ across all arrangements estimated by Fuller et al. (2014). Outlier genes within inverted regions have significantly lower mean estimates of ρ than non-outlier genes (Table 5 and Figure 8). These data show strongly differentiated genes interspersed among regions that experience higher levels of recombination within the inversion especially in the middle of the inversion which should be sufficient to decrease LD (Schaeffer & Anderson, 2005).

**Table 5.**
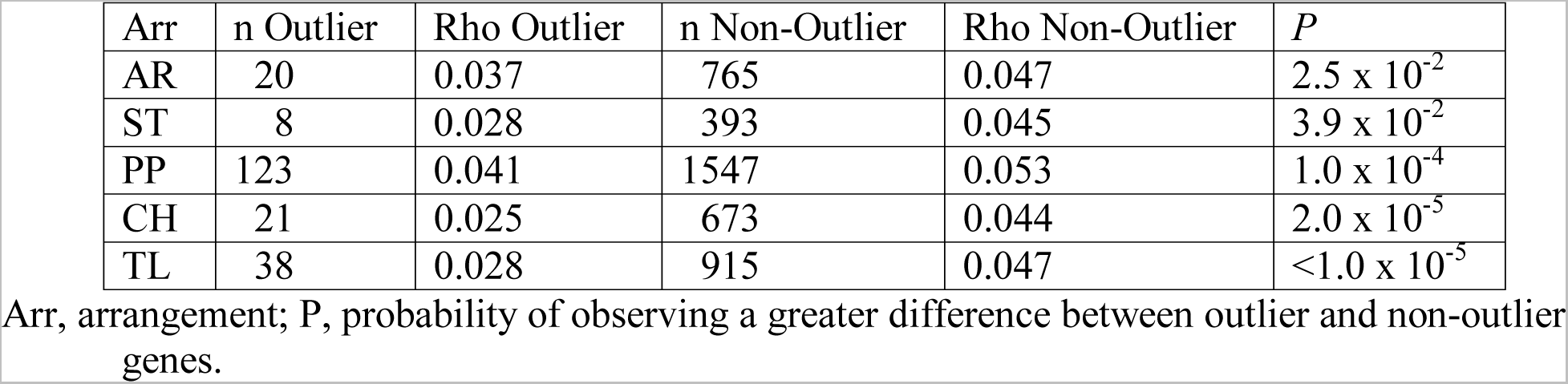
Mean estimates of the recombination parameter *Rho* in outlier and non-outlier genes within inverted regions.

| Arr | n Outlier | Rho Outlier | n Non-Outlier | Rho Non-Outlier | *P* |
| --- | --- | --- | --- | --- | --- |
| AR | 20 | 0.037 | 765 | 0.047 | $2.5 \times 10^{-2}$ |
| ST | 8 | 0.028 | 393 | 0.045 | $3.9 \times 10^{-2}$ |
| PP | 123 | 0.041 | 1547 | 0.053 | $1.0 \times 10^{-4}$ |
| CH | 21 | 0.025 | 673 | 0.044 | $2.0 \times 10^{-5}$ |
| TL | 38 | 0.028 | 915 | 0.047 | $<1.0 \times 10^{-5}$ |

Arr, arrangement; P, probability of observing a greater difference between outlier and non-outlier genes.

**Table 6.** GO analysis for genes in arrangements with significantly high mean DAF and PSBL and at least one fixed amino acid difference.

| Term | Function | Count | *P* Value | Fold Enrichment | Benjamini (*q*) |
| --- | --- | --- | --- | --- | --- |
| GO:0016021 | Integral component of membrane | 69 | 6.51E-06 | 1.47 | 9.77E-05 |
| GO:0050909 | Odorant binding | 11 | 1.09E-05 | 6.04 | 8.75E-04 |
| dpo04080 | Neuroactive ligand-receptor interaction | 6 | 3.15E-04 | 9.33 | 1.34E-03 |

**Figure 8.**
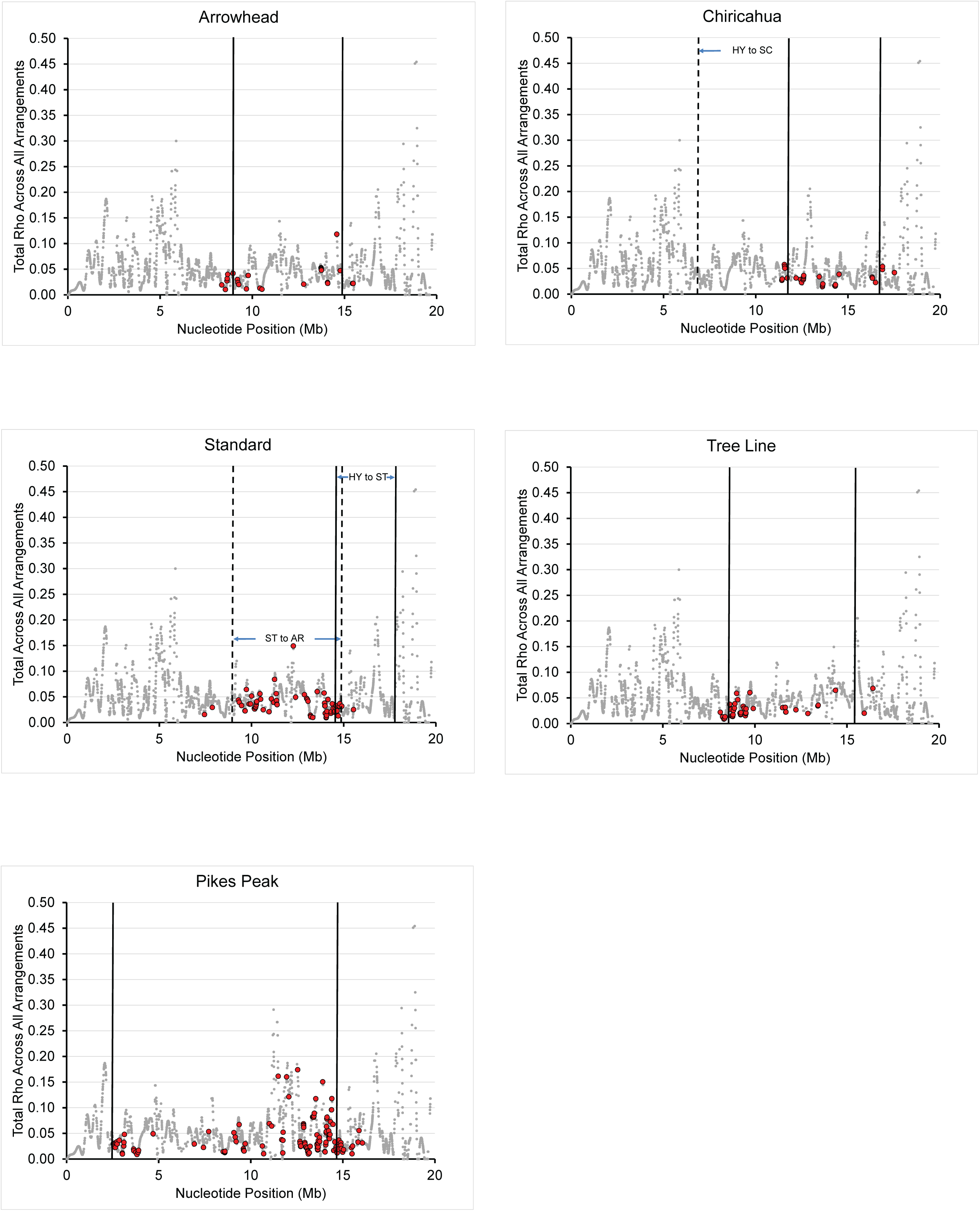
Estimates of the recombination parameter ρ within 2,669 gene regions across the third chromosome. Regions that showed a significantly large mean DAF or PSBL given the number of SNPs are indicated with a red circle. Regions that did not have an extreme mean DAF value are shown with a gray dot. The locations of inversion breakpoints are shown with a solid or dotted line. The order of genes is specific to the particular gene arrangement background.

Each arrangement has at least one gene with multiple fixed amino acid changes in our sample. Of these, the GA16823 gene in CH has the smallest number of fixed amino acid differences (9) while GA24454 in PP is the most extreme case with 47 fixed changes. In *D. melanogaster*, the ortholog of GA24454 is CG33017 and polymorphisms in the gene are significantly associated with olfactory responses to 2-phenyl ethyl alcohol (Arya et al., 2015). GA24454 is within inverted regions when PP and CU are paired with AR, ST, CH, or TL, however, the gene is outside inverted regions in heterozygotes in all other heterokaryotypes. Even though 207 amino acid polymorphisms are segregating in GA24454 across all arrangements, none of the other arrangements has a fixed amino acid difference. Coalescent simulations show that 47 fixed amino acid changes is greater than expected with a model of nested subsamples given 207 segregating amino acids (P=0.028; Hudson & Kaplan, 1986; Schaeffer et al., 2003). Furthermore, PP has only three shared amino acid polymorphisms while non-PP arrangements have 12 to 26 shared amino acid polymorphisms. The significantly large number of fixed amino acid differences in PP is evidence for adaptive evolution in GA24454.

### Lack of Extended Haplotype Homozygosity Surrounding Fixed Unique Nucleotide Alleles

At each fixed, derived site within an arrangement, we estimated the integrated haplotype score (*iHS*) to test for extended haplotype structure (See Figure S41 in the Supporting Information). Long stretches of extended homozygosity within an arrangement are likely to be associated with a strong or recent selection event. In each arrangement, we detected at least one site with an extreme positive or extreme negative (±3) *iHS* value. Non-neutral forces are expected to generate clusters of consecutive sites with extreme *iHS* values. We find evidence for only one such interval in any arrangement, an ∼18-kb stretch from 9,271,032 to 9,289,710 in AR. The interval intersects the coding regions of two overlapping genes (GA12653 and GA24326) neither of which showed a significantly elevated mean DAF or PSBL. For all other arrangements, no significant intervals of consecutive extreme *iHS* scores were observed.

### Third Chromosome Linkage Disequilibrium Patterns

Now we ask whether significant associations between sites decay with distance across the third chromosome (Figure 9). In *D. melanogaster*, LD tends to decay rapidly within the genome through recombination (Langley et al., 2012; Mackay et al., 2012), although recombination appears to be suppressed across the length of the inverted regions here (Fuller et al., 2016). In proximal and distal regions of the chromosome, we observe few pairwise comparisons that are in significant LD (3.97% of all comparisons). However, we find extensive LD generated within inverted regions (20.29% of all comparisons) and observe a significantly elevated proportion of significant pairwise comparisons present relative to non-inverted segments (Wilcoxon rank-sum test, *P*<0.05). Furthermore, multiple arrangements show significant associations between SNPs and arrangements with Fisher’s Exact Test across the majority of the chromosome (Figure 9) indicating that the pattern of LD is not being driven by a single arrangement. The exception is in the region where only PP differs from the other arrangements. Nucleotide variation around the proximal and distal breakpoints did not always show significant LD (Figure S42 in Supporting Information).

**Figure 9.**
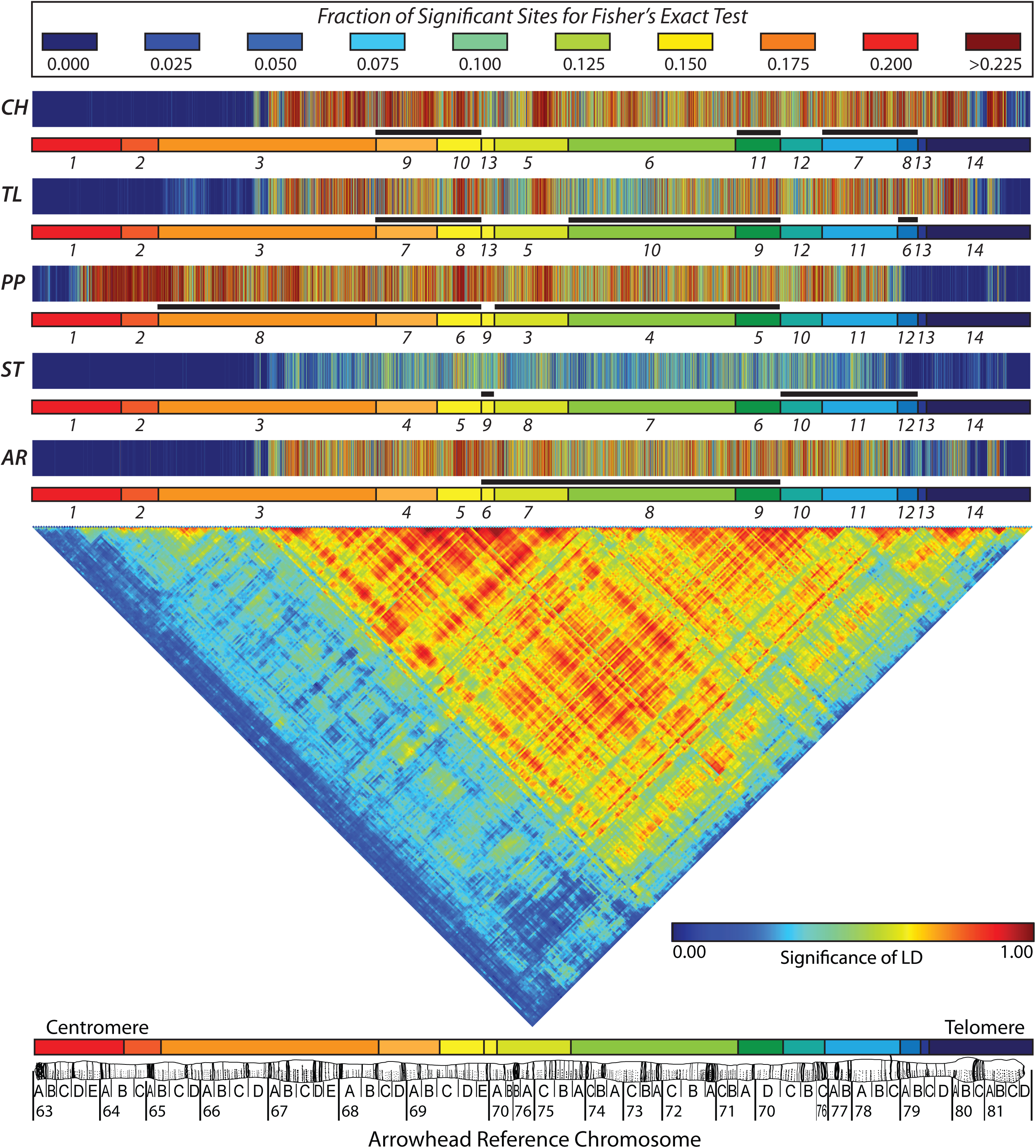
Triangular heat map showing the significance of linkage disequilibrium (LD) for all polymorphic sites on the third chromosome using the correlation-based procedure of Zaykin et al. (2008). Each point is calculated as the average of 100 adjacent sites. Red indicates greater LD and blue represent associations that are not significant. The top bars depict the results of a Fisher’s Exact Test for significant associations of alleles with a particular arrangement. The colors represent the proportions of significant tests (*p* < 1 x 10^-6^) in blocks of 100 sites.

### Evidence for Differentiation among Inversions, but not Populations

The six arrangements sampled were collected from five different geographic locations. We found minimal evidence for geographic differentiation (Table 2). Population-specific singletons account for 9.8 to 19.8% of the total unique polymorphism and the majority of derived polymorphisms (70.9 to 89.3 %) are shared among populations. The Mexican (SPE) population did have 56 (0.01%) unique fixed differences, but the other four populations did not have fixed unique derived polymorphisms. The extensive shared polymorphisms among geographic locations and lack of fixed differences are consistent with extensive gene flow among populations.

Gene arrangements, on the other hand, are differentiated from each other. Inversion specific polymorphism with a frequency of 1 in our sample account for 13.5 to 23.6 % of the observed unique polymorphism. Despite suppressed recombination in inverted regions, the majority of derived polymorphisms (56.3 to 78.7 %) are shared among arrangements. All arrangements have a greater proportion of fixed unique polymorphisms (0.8 to 7.4%) than the geographic populations. Gene arrangements harbor a substantial proportion of fixed unique sites, even in the presence of extensive shared polymorphism.

### Gene Ontology

To test for enrichment of common biological functions in the genes displaying evidence of adaptive evolution, we performed a Gene Ontology (GO) analysis using DAVID (v6.8) software (Huang, Sherman, & Lempicki, 2009a, 2009b). We note that direct experimental evidence is needed to confirm any of our results, however our analyses provide insight into genes with signals of adaptive evolution involved in similar biological functions that may underlie the targets of selection acting on the inversion polymorphism. For genes across all arrangements with at least one fixed amino acid change and significant mean DAF and PSBL, after correcting for multiple testing there is a significant enrichment of categories including odorant binding (*q* < 8.75 x 10^-4^; see Table 5) and neuroactive ligand-receptor interaction (*q* < 1.34 x 10^-3^). Several other categories of potential biological interest, including starch and sucrose metabolism and limonene and pinene degradation, contain genes which show evidence of adaptive evolution, although they are not significantly enriched.

Limonene and pinene are monoterpenes that are produced by a variety of plant species, including several coniferous pines, which have fungicidal properties and act as repellants to insects such as the mountain pine beetle (*Dendroctonus ponderosae*) (Wang et al., 2014). The composition of limonene and pinene in the resin of the Ponderosa pine (*Pinus ponderosa*) varies clinally across the Southwestern United States and in the overlapping species range of *D. pseudoobscura* (Smith, 1977). Furthermore, *D. ponderosae* has been shown to differentially colonize pines depending on the composition of monoterpenes, indicating that the species has evolved the ability to preferentially recognize the relative abundance of limonene and pinene (Thoss & Byers, 2006). A number of significant differentially expressed genes between *D. pseudoobscura* arrangements are also involved in limonene and pinene degradation (Fuller et al. 2016). Despite the wealth of genetic information supported by nearly a century of research in *D. pseudoobscura*, little is known regarding the species’ ecology and life history in nature. Our results here may suggest a link between the environment of *D. pseudoobscura* and genes showing evidence of adaptive evolution involved in sensory perception, metabolism and limonene and pinene degradation.

## Discussion

### Test of the Position Effect Hypothesis

The breakpoints examined here reject the position effect hypothesis in a narrow sense because none of the breaks disrupted the coding sequences of genes. Breakpoints may also generate position effects by altering the expression of boundary genes. Fuller et al. (2016) found only one case where a gene immediately adjacent to a breakpoint was differentially expressed. The gene GA22082, which is adjacent to the nearly coincident pHYST and dSTPP breakpoints, is expressed higher in ST than in PP chromosomes in larvae. Inversion events could also disrupt gene expression beyond breakpoints by altering topologically associated domains (TADs) (Hou, Li, Qin, & Corces, 2012). TADs partition the genome into structural domains within the nucleus that may be important for the coordination of gene expression. We inferred TADs in *D. pseudoobscura* based on synteny with TADs in *D. melanogaster* (Hou et al., 2012) (unpublished data). The 14 inversion breakpoints involve 20 *D. melanogaster* TADs. Five breakpoints (pHYSC, dSCCU, pSTAR, pSCTL, and dSTAR) disrupt a TAD and contain one or two differentially expressed genes (max: 2). The ST to AR event is particularly interesting because it splits a 79 kb TAD in half and leads to differential expression of two genes within the TAD on either side of the break. These results are intriguing, but should be viewed with caution because we do not have experimental evidence to suggest that TAD structure has been conserved between *D. melanogaster* and *D. pseudoobscura*.

There are other possible mechanisms for inversions to directly generate variation in the population. Novitski (1967) suggested that crossing over in inversion heterozygotes could set up conditions for meiotic drive to operate. Certain cross over events could lead to chromatids that differ in length such that shorter chromatids are transmitted at higher frequencies. Finally, Corbett-Detig (2016) showed that large inversions are often in close proximity to “sensitive sites” in the *D. melanogaster* genome. Sensitive sites normally promote crossing over, however, an inversion can lead reduce levels of genetic exchange when near the sensitive sites. Corbett-Detig (2016) concluded this can lead to a selective advantage for the inversion. Here, we conclude that breakpoint lesions have had minimal effects in generating phenotypic variation through the structural or expression alterations of boundary genes for selection to act on, however, further studies that map *D. pseudoobscura* TADs in different arrangements, test for meiotic drive in heterozygotes, and map sensitive sites in *D. pseudoobscura* are needed to fully reject the position effect hypothesis.

### Gene Phylogenies Support the Unique Origin of the Gene Arrangements

The phylogeny based on SNP variation across the third chromosome mirrors the history of the inversion mutations. Phylogenetic analysis supports the unique origin of the different arrangements except for regions near the centromere and telomere where recombination is prevalent. The expected phylogeny of the different arrangements is largely consistent with the cytogenetic phylogeny (Dobzhansky & Sturtevant, 1938), although some syntenic regions show an aberrant clustering of PP with TL.

One possible explanation for aberrant cluster of PP and TL is that there was an ancestral genetic exchange event. PP and TL have 739 SNPs that are shared between the arrangements in our samples. Although these regions are located within inverted regions of PP/TL heterozygotes, there is a 6.3 Mb region that can pair in the heterokaryotypes including regions (68C-69D, 69D-70A, 70B-70C, and 70D-74B). PP and TL do occur in the same populations and heterokaryotypes can form at appreciable frequencies allowing for recombinants. Indeed, PP/TL heterokaryotypes frequencies have been recorded as high as 32% in Mexico (Salceda, Guzman-Rincon, Rosa, & Olvera, 2015). The size of the region affected suggests that perhaps a double cross over event generated shared variation between the two arrangements (Arcadio Navarro et al., 1997) and the exchanged segment may have transferred adaptive genes based on a number of outlier genes that map to this segment.

### Evolutionary Forces for the Establishment and Maintenance of New Inversions

Here, we provide novel insights into the evolutionary forces that establish new inversion mutations in populations. If a new inversion increases by strictly neutral forces, as the arrangement increases in frequency, it will carry the allelic variants initially captured from the ancestral arrangement along with it and lead to extensive LD across the inverted region. Over time, LD of SNPs associated with the gene arrangement will decrease through genetic exchange except nearest to the breakpoints (Arcadio Navarro, Barbadilla, & Ruiz, 2000; Arcadio Navarro et al., 1997; Peischl, Koch, Guerrero, & Kirkpatrick, 2013). How much decay depends on the size of the inversion with larger inversions showing a greater reduction in LD in central regions than smaller inversions (Arcadio Navarro et al., 1997). The ages of the *D. pseudoobscura* arrangements are 0.5 to 1.4 million years old, which should be sufficient time for LD of variation to decay except near the breakpoints.

We can rule out genetic drift as an explanation for the inversion polymorphism in *D. pseudoobscura*. If genetic drift is responsible for the establishment and maintenance of the arrangements, their frequency is expected to be related to their relative ages (Kimura & Ohta, 1973), leading to higher levels of variation (Arcadio Navarro et al., 2000) and lower levels of LD (Hartl & Clark, 1997; Toomajian, Ajioka, Jorde, Kushner, & Kreitman, 2003). The strongest evidence against genetic drift is the pattern of variation in the AR chromosome. AR is the most frequent arrangement in the Southwestern United States suggesting that it is one of the oldest chromosomes, yet AR is relatively young because it was derived from the ST arrangement. This implies that AR emerged from the ST chromosome and has rapidly spread from California to Texas. The average value of Tajima’s *D*|*D*_min_| across AR is more negative than any other arrangement, which is consistent with a recent expansion. A similar observation is found in the closely related species *D. subobscura*, where allele frequencies of microsatellite markers have been shown to increase more than expected under genetic drift alone likely due to a recent expansion (J. Santos et al., 2016). The lack of significant windows of extended homozygosity in AR suggests that even though this arrangement is relatively young, there has been sufficient time for gene conversion break up LD. Additionally, genetic drift can be ruled out because each arrangement has multiple outlier genes which are spread across the inverted region. One would not expect outlier genes in the central region of the inversion because there has been sufficient time for gene conversion and crossovers to degrade LD.

Next, we can rule out a single beneficial sweeping allele because we find evidence for not one, but multiple outlier genes within the gene arrangements showing evidence of adaptive evolution. We find evidence of multiple genes with long branch lengths or elevated frequencies of derived mutations broken up by regions with high frequencies of shared polymorphisms. This is consistent with previous results showing multiple significantly differentially expressed genes between third chromosome arrangements maintained by the inversions (Fuller et al., 2016). Multiple loci are also thought to be functionally important for adaptive inversion clines in *D. melanogaster* (Kennington, Partridge, & Hoffmann, 2006).

The most likely explanation for the establishment and maintenance of chromosomal inversions in *D. pseudoobscura* is that inversions act as negative modifiers of recombination holding adaptive haplotypes together. By limiting genetic exchange, the original inversion mutation captures sets of linked alleles and maintains their associations. This is supported by the numbers of outlier genes that show evidence of adaptive evolution and the pattern of LD across the chromosome. Our surprising finding was that regions of the chromosome with high levels of LD are interspersed with regions of low LD or high values of ρ. In addition, we observed few regions of unusual haplotype structure or extended homozygosity within arrangements. We hypothesize that gene conversion and double crossovers can break up associations across an arrangement, but purifying selection removes any maladaptive combinations, consistent with inversion establishment models (Charlesworth & Charlesworth, 1973). These results should be interpreted with caution because some of the shared polymorphism we observe may be due to gene conversion events from balancer chromosomes used to create the isogenic strains (Miller et al., 2016).

Two models are consistent with the establishment of inversions to reduce recombination, the local adaptation and epistasis models, which may not be mutually exclusive. The distinction between these two hypotheses is whether the gene products interact at the molecular level or have a synergistic effect on fitness. Under the local adaptation model, there is no need for the gene products to interact or act synergistically either in a genetic pathway or as transcription factors regulating a common gene (Kirkpatrick & Barton, 2006). The geographic range of *D. pseudoobscura* straddles several physiographic provinces (Lobeck, 1948) and adult flies are capable of dispersing among these ecological niches. Numerical analysis has shown that geographic inversion frequencies of *D. pseudoobscura* are consistent with a model of selection in heterogeneous environments (Schaeffer, 2008). However, it remains unclear why multiple arrangements exist in single populations under the local adaptation model and further work is needed to characterize the local fitness environments in ecological niches

The data presented here supports the hypothesis that the size of the inversion may be an indirectly selected character (Caceres, Barbadilla, & Ruiz, 1997, 1999). The ST arrangement is an exceptional case because the majority of the candidate selected genes are outside of the original derived inversion (HY to ST) and instead found in the proximal region, which overlaps with the AR inversion. Because ST gave rise to AR and AR remained in populations with ST, it appears that new adaptive genes in ST have become fixed from standing variation after the HY to ST inversion event. We hypothesize this resulted from selection on standing variation within ST (Wyatt W. Anderson et al., 1991; Dobzhansky, 1944) and suppression of recombination due to the presence of AR and CH in populations with ST. Thus, our data suggest that an initial inversion event can lock up adaptive combinations of genes within the inverted segment, but additional multi-locus combinations can be established outside of the initial inversion breakpoints based on what other arrangements are present in the population.

If double cross overs are responsible for breaking up LD, one would expect larger regions of sharing between arrangements (Arcadio Navarro et al., 1997), especially between ST and PP. The ST to PP inversion corresponds to 52% of the length of the third chromosome and one would expect seven percent of gametes from a ST/PP heterozygote to produce double cross overs leading to more similarity within their inverted regions. Homogenization, however, is unlikely because ST and PP do not co-occur at appreciable frequencies in nature. Wallace (1953) proposed the triad model where coadapted gene complexes could be eroded when sets of arrangements separated by single inversion steps such as AR-ST-PP are present in the same population. Therefore, it is reasonable to speculate that all members of a triad do not co-occur for this reason. ST and PP support the triad model because of the non-trivial potential to generate maladaptive recombinant haplotypes.

There is a significant enrichment of genes involved in sensory perception and odorant binding showing evidence of adaptive evolution. Analyses of insect genomes have found that odorant perception and detoxification proteins are typically found to be amplified in copy numbers and show evidence of recent positive selection (Chen et al., 2015; Nene et al., 2007; Scott et al., 2014; Smadja, Shi, Butlin, & Robertson, 2009; The Honeybee Genome Sequencing Consortium, 2006; The International Aphid Genomics Consortium, 2010; Tribolium Genome Sequencing Consortium, 2008; You et al., 2013). Our study further suggests these genes may be targets of selection in insect species because they can play a role in adapting to complex environments. Furthermore, several of the genes showing evidence of adaptive evolution are members of the limonene and pinene degradation pathway. It is intriguing to speculate that ponderosa pine is part of the ecology of *D. pseudoobscura* either because the adults are associated with bark of the tree or larvae feed on the pine nuts.

The establishment of inversion clines may play a role in the speciation process. In such a system, the emergence of Dobzhansky-Muller incompatibility genes within one arrangement background can easily spread within the inversion type, but not to other chromosomal backgrounds (A. Navarro & Barton, 2003). This may have been how *D. persimilis* was formed (Noor, Grams, Bertucci, & Reiland, 2001). The distribution of *D. persimilis* is sympatric with *D. pseudoobscura* and the two species differ by several fixed inversion differences that include reproductive isolation genes. Thus, *D. pseudoobscura* may be ripe for additional speciation events.

The evolution of different gene arrangements in *D. pseudoobscura* may be in response to life history and environmental challenges that require many genes of small effect to adapt to a heterogeneous environment. Independent assortment and recombination play a vital role in helping an organism to generate multiple combinations for selection to sift through. While this is an advantage, it is also a curse because adaptive combinations of genes can be split apart as quickly as they are made. In this system, it seems that chromosomal inversions help to hold adaptive combinations together.

## Acknowledgements

Ian S. Leopold for preparation of polytene chromosome squashes, digital photography of the chromosomes, and assembly of the mosaic images of the chromosomes. Andre G. Wallace for sequencing the 18 regions used to validate the Illumina sequences. Richard Kliman at Cedar Crest College who kindly provided the SNP data for the *D. pseudoobscura* second chromosome. The authors would like to thank the staff at the Baylor College of Medicine Human Genome Sequencing Center for their technical and computational contributions. This work was supported by a grant from the National Institute for General Medical Sciences at the National Institutes of Health R01 GM098478 to S.W.S. and the Penn State-National Institutes of Health funded Computation, Bioinformatics and Statistics (CBIOS) Predoctoral Training Program to Z.L.F.

## Data Accessibility

DNA Sequences: NCBI SRA: SRX204748-SRX204792, SRX091323, SRX091311, SRX815755, SRX2484948, SRX2484950, SRX2484953, SRX2484955, SRX2484969.

SNP table for the third chromosome containing the sites used for all subsequent analyses is available through Scholarsphere (https://scholarsphere.psu.edu).

Computer Code is available: (https://scholarsphere.psu.edu)

## Author Contributions

S.W.S conceived the project. S.R., generated the sequence data. Z.L.F., G.D.H., S.R.and S.W.S, analyzed the data. Z.L.F., G.D.H., S.R., and S.W.S. wrote the manuscript.

